# Predicting reproductive phenology of wind-pollinated trees via PlanetScope time series

**DOI:** 10.1101/2024.08.02.606226

**Authors:** Yiluan Song, Daniel S.W. Katz, Zhe Zhu, Claudie Beaulieu, Kai Zhu

## Abstract

Airborne pollen triggers allergic reactions which result in public health consequences. A better understanding of flowering and pollen phenology could improve airborne pollen predictions and reduce pollen exposure. Data on the timing of flowering and pollen release are needed to improve models of airborne pollen concentrations, but existing *in-situ* data collection efforts are expensive and spatially sparse. Satellite-based estimates of plant phenology could potentially enable large-scale data collection, but it is difficult to detect the reproductive phenology of wind-pollinated flowers from space. Here, we infer the reproductive phenology of wind-pollinated plants using *PlanetScope* time series with a spatial resolution of 3 m and a daily revisit cycle, complemented by *in-situ* flower and pollen observations, leveraging the correlation between vegetative and reproductive phenology. On the individual tree level, we extracted PlanetScope-derived green-up time and validated its correlation to flowering time using flower observations in a national-scale observatory network. Scaling up to the city level, we developed a novel approach to characterize pollen phenology from PlanetScope-derived vegetative phenology, by optimizing two tuning parameters: the threshold of green-up or green-down and the time lag between green-up/down and flowering. We applied this method to seven cities in the US and seven key wind-pollinated tree genera, calibrated by measurements of airborne pollen concentrations. Our method characterized pollen phenology accurately, not only in-sample (Spearman correlation: 0.751, nRMSE: 13.5%) but also out-of-sample (Spearman correlation: 0.691, nRMSE: 14.5%). Using the calibrated model, we further mapped the pollen phenology landscape within cities, showing intra-urban heterogeneity. Using high spatiotemporal resolution remote sensing, our novel approach enables us to infer the flowering and pollen phenology of wind-pollinated plant taxa on a large scale and a fine resolution, including areas with limited prior *in-situ* flower and pollen observations. The use of PlanetScope time series therefore holds promise for developing process-based pollen models and targeted public health strategies to mitigate the impact of allergenic pollen exposure.

## 1 Introduction

Pollen is a trigger of allergic asthma and allergic rhinitis (hay fever), imposing significant costs on public health (Reid & Gamble, 2009; D’Amato et al., 2020; Idrose et al., 2022; A. B. Singh & Kumar, 2022). The onset, duration, and intensity of pollen seasons are highly related to the phenology, the timing of recurring biological events, of wind-pollinated plants. Health risks from pollen exposure are likely to exacerbate under global change, reflected in earlier starts and often longer durations of flowering seasons (Mo et al., 2017) and pollen seasons (Anderegg et al., 2021; Ziska et al., 2011, 2019), as well as higher pollen concentrations (Ziska & Caulfield, 2000). Currently, the inadequacy of pollen monitoring and publicly available pollen models hinder accurate assessments and timely public health responses to changing pollen seasons exacerbated by global changes. The elevating risks call for major improvements in the collection of flower and pollen phenology data on large scales and fine resolutions, in order to enhance mechanistic understanding and prediction of the reproductive phenology of wind-pollinated plants.

Previous research on plant reproductive phenology for public health has been limited by insufficient observational data. In the field of aerobiology, pollen phenology is studied with airborne pollen concentration data from air sampling, generally through national pollen monitoring networks (Scheifinger et al., 2013). These pollen concentration data are collected with systematic protocols at a daily temporal resolution, thus serving as a high-quality source for pollen modeling. However, air sampling of pollen is expensive, and the stations are spatially sparse (Anderegg et al., 2021). Aggregated over a large area and identified to the family or genus level, pollen samples often do not allow us to study fine-scale spatial variations and intra-genus variations. In the field of ecology, ground observations of flowering phenology, from observatory networks and community science, have been correlated with pollen phenology (Crimmins et al., 2017; Elmendorf et al., 2016; Templ et al., 2018). These phenological observations have a larger spatial coverage and a finer taxonomic resolution compared to air sampling data but are limited by subjectivity in the classification of phenophases (Donnelly et al., 2022) and spatiotemporal sampling bias (Pearse et al., 2017).

Although both air samples and ground observations have been used to advance pollen modeling, these models still need to be improved in accuracy and spatial robustness (Scheifinger et al., 2013). On the one hand, data-driven pollen models using statistics (Frenguelli et al., 1989) or machine learning (Seo et al., 2019; Zewdie et al., 2019; F. Lo et al., 2021) are usually site-specific and sometimes lack accuracy (Chuine & Belmonte, 2004; Maya-Manzano et al., 2021). It is therefore challenging to extrapolate locally-trained pollen models to locations without prior *in-situ* data collection. Integration of land surface phenology as predictors has been suggested to improve data-driven models (Huete et al., 2019; F. Lo et al., 2021). On the other hand, process-based pollen models that explicitly account for plant reproductive phenology, pollen production, and pollen dispersion have been shown to be promising in predicting pollen seasons with robustness across Europe (Chuine & Belmonte, 2004; Sofiev et al., 2006; Vogel et al., 2008; García-Mozo et al., 2009; Sofiev et al., 2015) and predicting spatial variations of pollen concentrations within cities in the US (Katz, Baptist, et al., 2023). Several gaps exist in the current process-based pollen models: they are rarely updated with near real-time observations; they are not available on an operational scale outside of Europe; they are usually limited in spatial resolution, missing important heterogeneity within cities (Katz & Batterman, 2020). To create generalizable and granular process-guided models of airborne pollen, we need to go beyond existing empirical data and obtain plant reproductive phenology data with a large spatial coverage and fine spatial resolution.

To overcome the data challenge, remote sensing has been explored to inform pollen and flower phenology, building on the correlation between reproductive phenology and vegetative phenology. Leaf out and flowering, are tightly linked phenological events in a plant’s life cycle, evolved to occur in a predictable sequence with stable time intervals (Davies et al., 2013; Guo et al., 2023). Such flower-leaf sequences are crucial to plant fitness in temperature regions (Buonaiuto & Wolkovich, 2021; Guo et al., 2023), such as through effective wind pollination in flowering-first species (Buonaiuto et al., 2021). Such a biophysical relationship motivates the inference of reproductive phenology from remotely-sensed vegetative phenology (Davies et al., 2013). For example, the onset of bud burst detected from Moderate Resolution Imaging Spectroradiometer (MODIS) Normalized Difference Vegetation Index (NDVI) and Global Inventory Monitoring and Modeling System (GIMMS) NDVI were found to correlate with the onset of birch flowering (Karlsen et al., 2008) and birch pollen season (Høgda et al., 2002), respectively, on continental and decadal scales. Interannual variations in the flowering time of multiple plant functional types have been explained by remotely sensed green-up time (Delbart et al., 2015). Moving beyond correlation, data-driven predictive pollen models have also benefited from incorporating MODIS Enhanced Vegetation Index (EVI) as a predictor (Huete et al., 2019; F. A. Lo, 2020; Yang et al., 2022). These studies show the feasibility of using satellite remote sensing to greatly expand the spatial coverage of reproductive phenology data and to improve pollen models.

Despite their broad spatial coverage, satellite remote sensing data products previously used to study flowering and pollen phenology have a limited spatial resolution, specifically 250 m (MODIS), 500 m (MODIS), or 8 km (GIMMS). Land surface phenology detected on this resolution suffers from the mixed pixel problem (X. Chen et al., 2018). This is particularly problematic for urban landscapes that are highly heterogeneous in land cover and plant species. Given that pollen exposure and plant reproductive phenology are highly spatially heterogeneous within a city (Katz et al., 2019; Katz & Carey, 2014), land surface phenology at a coarse spatial resolution does not satisfy the need for spatially-explicit pollen modeling for public health.

With a spatial resolution of 3 m and a daily revisit cycle, PlanetScope data provide an excellent opportunity to gather plant reproductive phenology data on a large scale and at an individual tree level. We identified the untapped potential of PlanetScope data for public health from two streams of research. On the one hand, PlanetScope data have been used to successfully detect large and brightly-colored flowers within a stand, with indices designed to capture spectral signatures of flowers, such as enhanced bloom index (EBI) (Campbell & Fearns, 2018; B. Chen et al., 2019; Dixon et al., 2021). Although supporting the use of PlanetScope to detect canopy-level phenological variations, the PlanetScope-derived bloom index can hardly be applied to wind-pollinated flowers that are small and inconspicuous (Kim et al., 2020).

Reproductive phenology of wind-pollinated flowers will therefore largely rely on the inference from vegetative phenology. On the other hand, PlanetScope-derived EVI has been shown to be a reliable data source for tree-level vegetative phenology, validated by other remote sensing data products (Moon et al., 2021) and ground observations (Moon et al., 2022; Zhao et al., 2022; Y. Liu et al., 2024). Despite promising applications of PlanetScope data to detect spatial variations among individual tree canopies and to derive vegetative phenology, it has not yet been used to infer tree-level flowering phenology from vegetative phenology. Further, to our knowledge, there has not been research linking PlanetScope directly to pollen phenology, which is central to modeling pollen exposure.

In this study, we assessed the potential of using vegetative phenology data extracted from PlanetScope to infer the flowering and pollen phenology of wind-pollinated trees. Specifically, the study focused on answering the following research questions (Fig. 1, Fig. S1).

**Figure 1.**
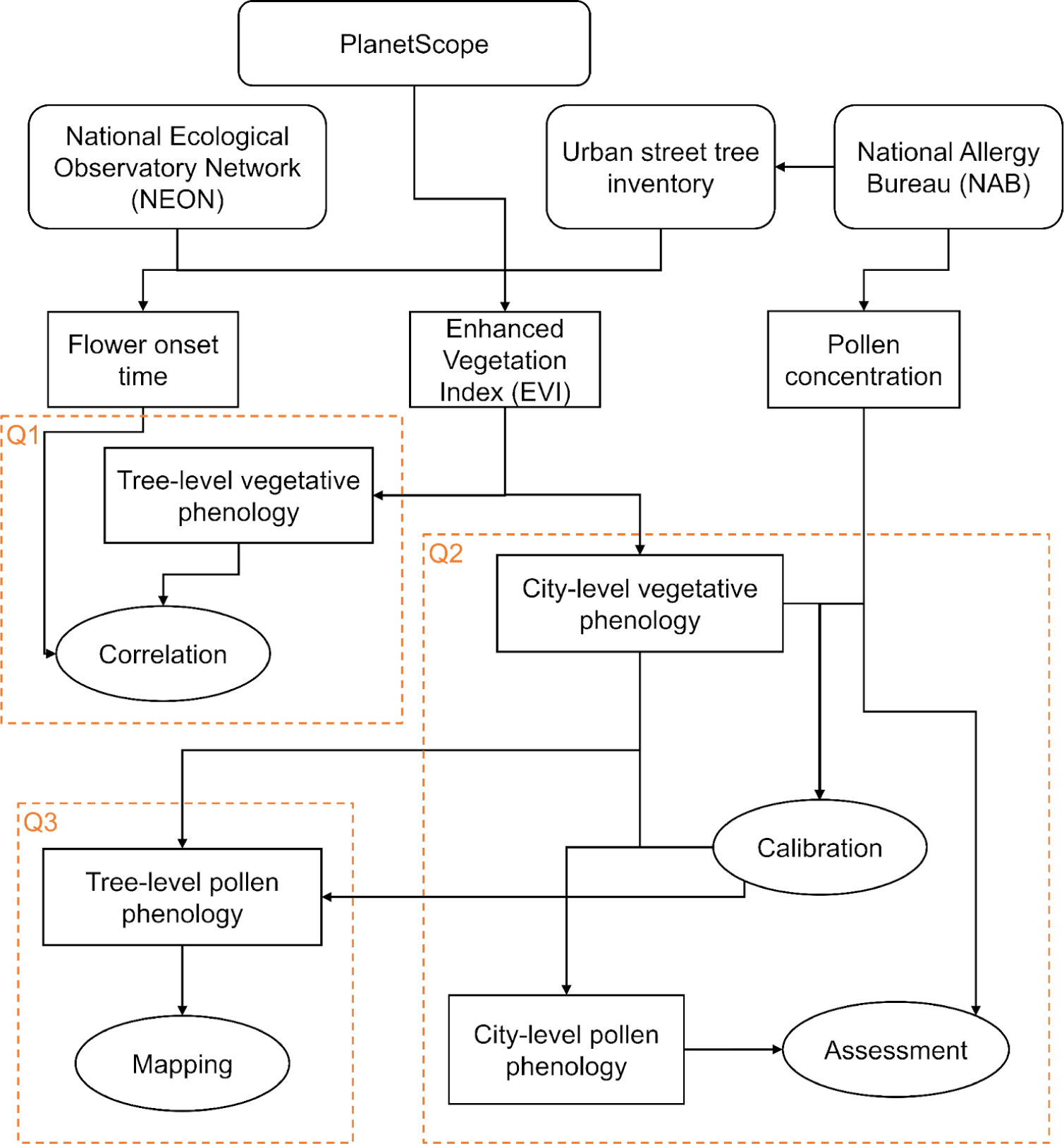
Simplified flow diagram of data and methods used in this study, and corresponding research questions. Rounded rectangles represent source datasets. Rectangles represent derived variables to analyze. Ellipses represent data analysis steps.

**Q1:** On the tree level, does PlanetScope-derived vegetative phenology correlate with flowering phenology monitored by field observations?

**Q2:** Upscaled from the tree level to the city level, can PlanetScope-derived vegetative phenology be used to accurately infer pollen phenology characterized by airborne pollen concentrations? In particular, does this inference extrapolate over a large spatial scale, to locations where airborne pollen concentration data are unavailable?

**Q3:** Scaling back to the tree level, can PlanetScope-derived vegetative phenology be used to explore fine-scale intra-urban heterogeneity of pollen phenology, i.e., a pollen allergy landscape?

In answering these two questions, we developed a novel workflow for obtaining cross-scale reproductive phenology data from PlanetScope time series. Our workflow has two core ideas: upscaling tree-level green-up/down time to the city-level green-up/down phenology, and tuning two parameters to calibrate city-level green-up/down phenology with city-level *in-situ* pollen phenology.

## 2 Materials and Methods

### 2.1 Data description

#### 2.1.1 Tree-level flowering observations

To test whether PlanetScope can capture tree-level variations in flowering phenology, we retrieved plant phenology observations from the National Ecological Observatory Network (NEON) (DP1.10055.001) that were integrated into the USA National Phenology Network (USA-NPN) (Elmendorf et al., 2016; Crimmins et al., 2017; National Ecological Observatory Network, 2020). At each site and every year, 90–100 tagged individual plants were observed *in situ* by trained technicians for their vegetative and reproductive phenophase status with varying sampling frequencies up to three times per week, following the phenophase definitions and protocols of NPN (Denny et al., 2014). We downloaded individual phenometrics from the NEON data submitted to NPN, which are the estimates of the dates of phenophase onsets and ends, measured from a series of consecutive “yes” phenophase status records. In this study, we used flower and leaf onset dates, which are the time of first “yes” observations for an individual tree in a given year. To complement the phenological data, we retrieved the accurate coordinates of tagged NEON trees using the R package *geoNEON* (National Ecological Observatory Network, 2023). We focused on primary sampling sites within the continental United States (CONUS) that have available coordinates of tagged individual plants (Fig. S2). We included data from 2018 to 2022, as fully operational PlanetScope data collection started in 2018 (Fig. S3).

We focused on deciduous wind-pollinated tree species with considerable public health impacts in the CONUS and Southern Canada region (F. Lo et al., 2019). They include seven genera ranked by percent abundance in pollen-producing taxa: *Quercus* spp. (oak), *Morus* spp. (mulberry), *Ulmus* spp. (elm), *Fraxinus* spp. (ash), *Betula* spp. (birch), *Acer* spp. (maple), and *Populus* spp. (poplar, aspen, cottonwood). *Ulmus* spp. were considered to have an early- and a late-flowering group, whose phenology was analyzed separately.

#### 2.1.2 City-level pollen concentration from air sampling and street-tree inventory

To examine the potential of using PlanetScope for city-level pollen phenology and to inform public health, we obtained consistent and accurate pollen concentration data from stations associated with the American Academy of Allergy, Asthma & Immunology (AAAAI) National Allergy Bureau (NAB) (AAAAI, 2022) (Figs. 2, 3). At around 80 stations located throughout the continental US, airborne pollen was sampled daily using volumetric impactor samplers (mostly Burkard samplers) (Portnoy et al., 2004), and then classified to the genus or family level and counted by NAB-certified operators. The NAB dataset is the most commonly used data source for the description and prediction of pollen phenology in public health and ecological research in North America. We obtained pollen concentration data collected until late 2023 (Fig. 3). Data were available during most springs and summers after the establishment of sampling stations, with missing data when airborne concentrations were low (Fig. S4). As in the tree-level analysis, we focused on the pollen phenology of seven wind-pollinated tree genera (*Quercus*, *Morus*, *Ulmus*, *Fraxinus*, *Betula*, *Acer*, and *Populus*).

**Figure 2.**
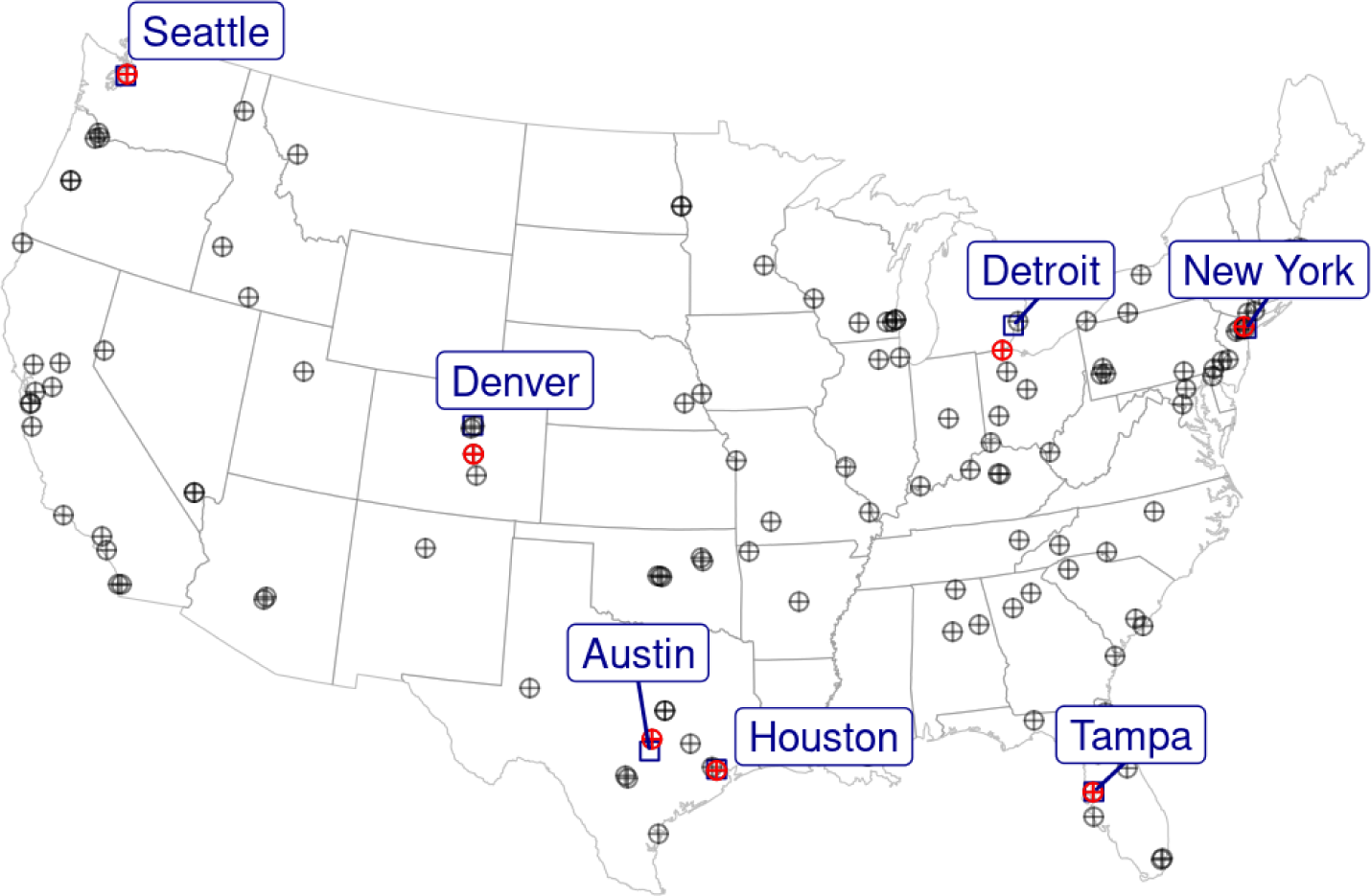
Map of seven studied cities with street tree inventory (blue squares) and pollen monitoring stations associated with the National Allergy Bureau (NAB) (red crossed circles). Pollen monitoring stations were located within 100 km of the centroid of the city’s street trees. Other NAB pollen monitoring stations not used in this analysis are marked in gray crossed circles.

**Figure 3.**
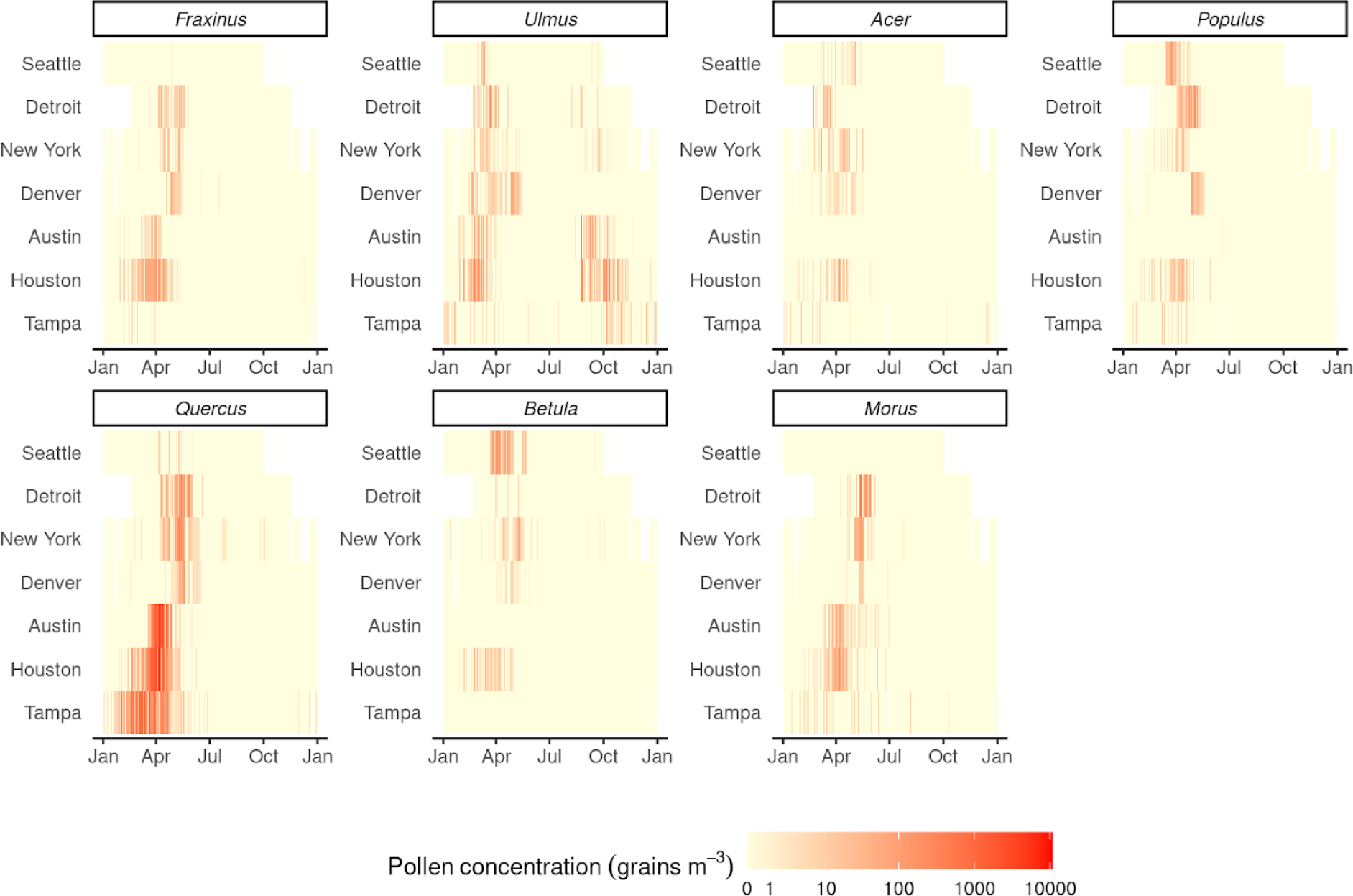
Daily climatology of pollen concentration (grains m^-3^) of seven key pollen-producing genera in studied cities. Climatologies are based on averages over the period of 2003–2023. Cities are ordered according to their latitude. Data are from the National Allergy Bureau (NAB) pollen monitoring stations.

To complement the pollen concentration data, we obtained street tree inventories in selected cities (Figs. 2, 4; details on sources of tree inventories in Table S1). Necessary reprojections were performed to convert all coordinates in street tree inventories to longitude and latitude. The taxonomy of street trees was resolved with the R package *taxize* (Chamberlain & Szöcs, 2013) for selecting trees in the genus of interest. When there were more than 2,000 recorded trees of a genus in a city, we randomly selected 2,000 trees (Psutka & Psutka, 2019).

**Figure 4.**
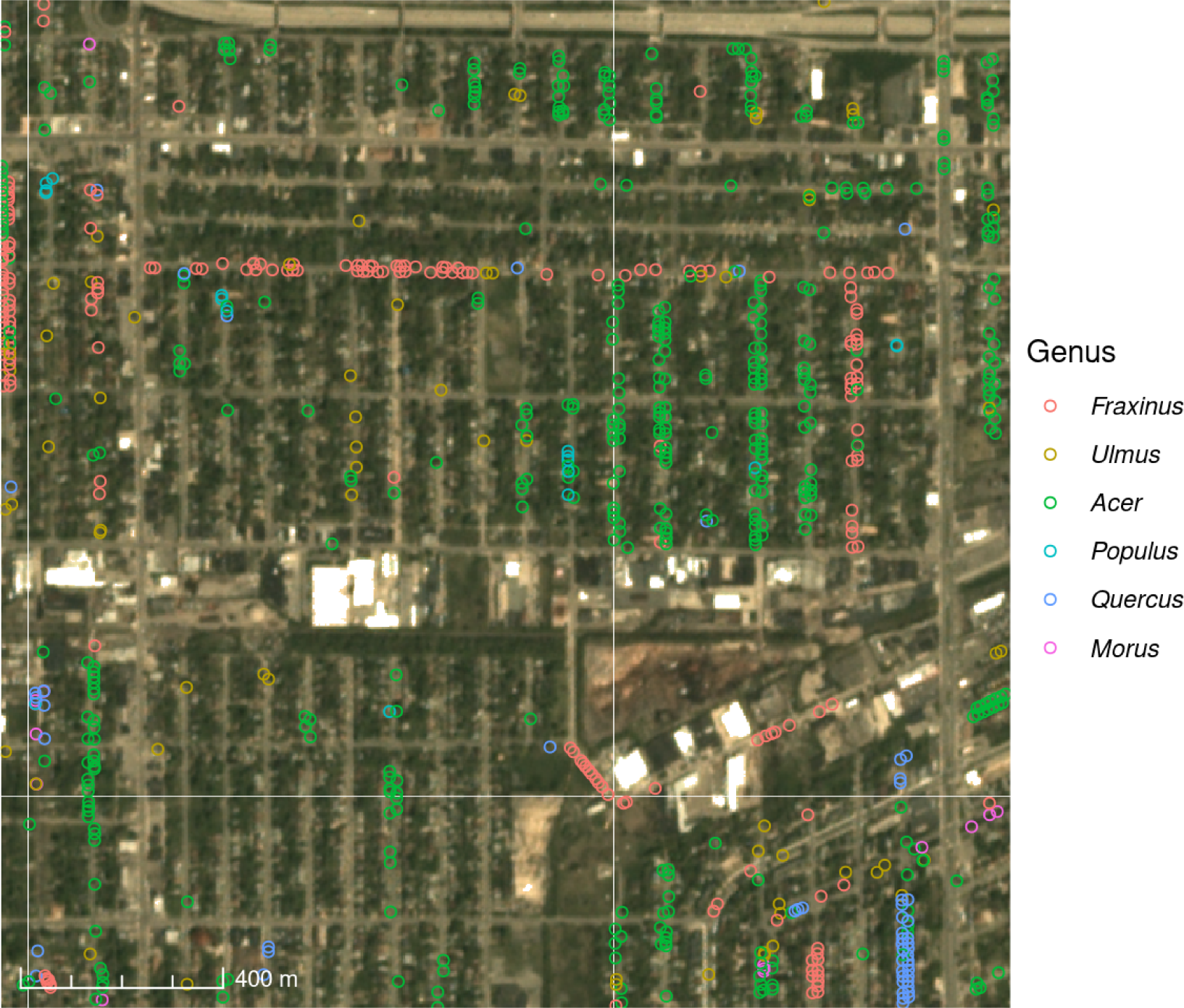
A subset of street trees in Detroit overlayed on a true-color PlanetScope image on May 8, 2017. The colors of points indicate different genera.

We focused on seven US cities with an available street tree inventory and a nearby pollen monitoring station (Fig. 2): Austin (AT), Detroit (DT), Denver (DV), Houston (HT), New York (NY), Seattle (ST), and Tampa (TP). Most focal cities have a pollen monitoring station within the city, with a mean distance from the pollen monitoring station to the centroid of all censused street trees ranging from 3.9 km to 41 km. Two exceptions were that Denver’s pollen concentration data were from Colorado Springs (mean distance 97 km) and that Detroit’s pollen concentration data were from Sylvania (mean distance 91 km).

#### 2.1.3. PlanetScope reflectances for vegetative phenology

We retrieved PlanetScope images from 2017 to 2023 for all trees involved in the analyses, including sampled trees at NEON sites and street trees from seven wind-pollinated tree genera in seven selected US cities (Fig. 4). We downloaded the PlanetScope atmospherically corrected surface reflectance product (ortho_analytic_4b_sr) (Planet Team, 2017) through the Planet API, using a custom package based on the R package *planetR* (Bevington et al., 2019/2024). We applied the “harmonize” tool in the Planet API with “Sentinel-2” as the target sensor, in order to make all PlanetScope data consistent and approximately comparable to Sentinel-2 data (Kington & Collison, 2022). All images obtained were acquired during the day (sun elevation > 0 m). At the coordinates of the trees of interest, we obtained the reflectances in the red, green, blue, and near-infrared bands. For quality control, we applied Usable Data Masks (UDM2) (Planet Team, 2023) to include only pixels that were clear, had no snow, ice, shadow, haze, or cloud, and had algorithmic confidence in classification ≥ 80%. Given little information on the size and shape of trees, we obtained reflectances at the focal coordinates, instead of within polygons that cover tree canopies (Dixon et al., 2023). For each date, pixel, and band, we computed mean reflectances if there were multiple visits in a day.

### 2.2 Data processing

#### 2.2.1 Curating ground-observed flowering phenology data

In processing the NEON flowering phenology data, we exclude NEON sites located outside the continental US. We also excluded sites situated in the Mediterranean climatic zone (all sites in California), where plant phenology is primarily driven by precipitation instead of temperature, distinct from other parts of the continental US. We focused on wind-pollinated tree species that were widely represented in the NEON data (≥ 50 records). We removed outliers of spring flower onset dates that were biologically implausible for the species present in the dataset (later than day 150).

#### 2.2.2 Curating air-sampled pollen concentration data

We processed NAB pollen concentration data to characterize pollen phenology in several steps:

1. In order to include at least one full pollen peak, we extended data in each year from day-of-year (DOY) 275 (Oct 2) in the previous calendar year to day 90 (Mar 30) in the following year. Data on day 366 in leap years were ignored.
2. We removed combinations of genus and city when there were no trees of interest in the street tree inventory, or there were no more than 30 pollen concentration records greater than or equal to 0 grains m^-3^. We removed *Fraxinus* spp. from New York and Detroit from our dataset due to the mass die-off of these trees in these two cities during our study period.
3. To compress extreme values and stabilize the variance, we transformed all pollen concentration values 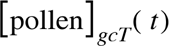 to their square root 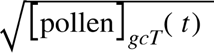 (Bastl et al., 2018; Bonini et al., 2022). Here, *t* represents the day of year. Indices *g*, *c*, *T* represent genus, city, and year, respectively.
4. In order to focus on single pollen peaks for plant genera that have both early- and late-flowering variations (e.g., *Ulmus* spp.), as well as to reduce the confounding effect of outliers outside the reproductive season, we constrained the pollen seasons for each taxon, setting the pollen concentration outside the season to zero. Genus-specific pollen seasons were determined by summing the total pollen concentration over all cities and years, fitting a Gaussian kernel, calculating a window of mean ± 1.96×standard deviation (Zhang & Steiner, 2022), and extending the window by 50 days on both ends (Fig. S5). An exception was that the early and late pollen windows of *Ulmus* spp. were detected by fitting a Gaussian mixture model with two peaks.

To handle short gaps of missing data within the pollen peaks and reduce the impacts of outliers, we gap-filled and smoothed the time series with weighted Whittaker smoothing 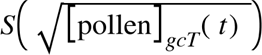 (Eilers, 2003, 2004). Here *S* refers to a weighted Whittaker smoothing operation.

#### 2.2.3 Calculating Enhanced Vegetative Index from PlanetScope reflectances

In order to characterize tree-level vegetative phenology, we used reflectances from PlanetScope images to extract phenological metrics, specifically green-up or green-down time, for each individual tree of interest. We performed the following steps.

1. We calculated the enhanced vegetation index (EVI) (H. Q. Liu & Huete, 1995) (Eqn. 1).

PlanetScope EVI has been shown to accurately extract leaf phenology metrics validated by local digital camera imagery (PhenoCams), robust to different atmospheric conditions and less likely to saturate in densely vegetated areas compared to PlanetScope normalized difference vegetation index (NDVI) (Wu et al., 2021).

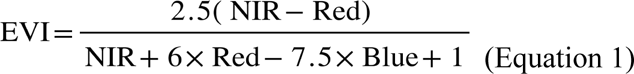

We used the following criteria to filter out possibly erroneous EVI values: reflectances in all visible bands were positive values, and EVI was between zero and one.

1. We extended the time series in each year from day 275 (Oct 2) in the previous calendar year to day 90 (Mar 30) in the following year in order to include at least one full growing season with green-up and green-down. This step was necessary for the detection of green-up day when EVI increases from the minimum before the New Year, and the detection of green-down day when EVI decreases to the minimum after the New Year. We gap-filled and smoothed all extended time series with weighted Whittaker smoothing (Kong et al., 2019).
2. We selected a time series of EVI with significant seasonality. In particular, we fitted a simple linear regression model and then three piecewise regression models with one, two, and three change points, respectively (Eqn. 2) (Beaulieu & Killick, 2018).

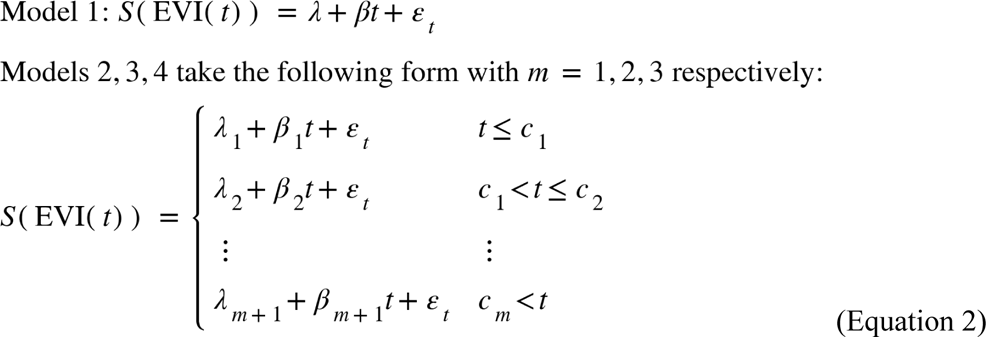

EVI time series from each genus, city, year, and tree were analyzed separately, but we omit the indices here for simplicity. In Model 1, *t* represents the time in days, *λ* and *β* represent the intercept and trend and *ε_t_* is white noise. In Models 2-4, *m* represents the number of changepoints, *c_m_* (*m* = 1,..,3) represent the timing of change points, and *λ*_1_,…,*λ_m_* and *β*_1_,…,*β_m_* represent the intercept and trend in each segment. Piecewise regression models were fitted with R package *segmented* (Muggeo, 2008). We ranked the four models according to the Akaike information criterion (AIC). If a simple linear regression was the best model, we discarded the time series as it may lack seasonal changes in greenness.

### 2.3 Inferring reproductive phenology with PlanetScope

#### 2.3.1 Inferring flowering phenology on the tree level

For individual trees at NEON sites (Fig. S1) monitored for phenology, we used the EVI time series to identify the green-up phases empirically (Fig. 5). The end of a green-up phase was determined as the day of year when EVI reaches the maximum in the growing season. The start of a green-up phase was then determined as the day of year when EVI is at the minimum, prior to the end of the green-up phase. We then determined the timing of green-up at the 50% threshold.

**Figure 5.**
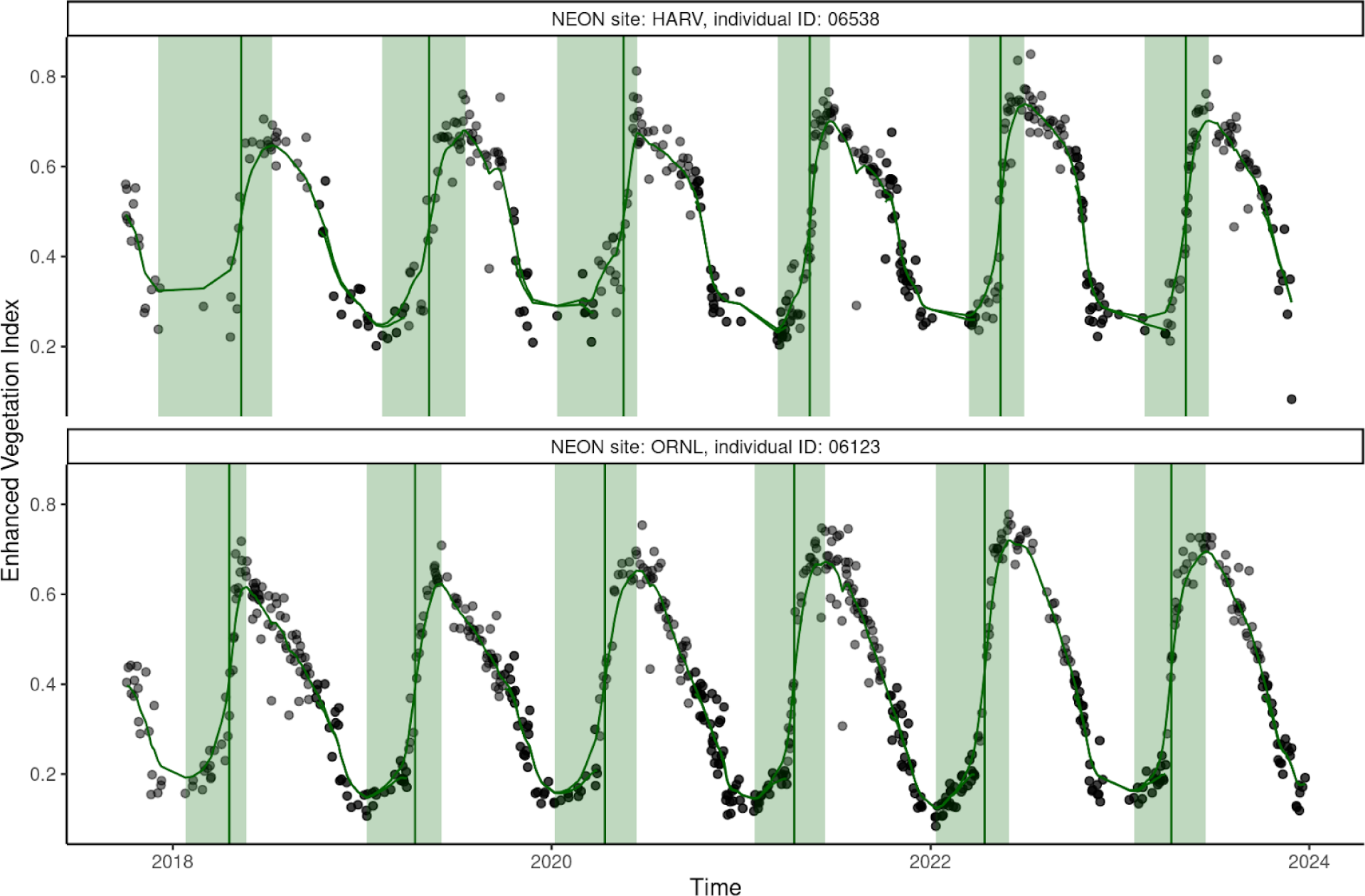
Extraction of tree-level phenological metric from PlanetScope data for wind-pollinated trees sampled at the National Ecological Observatory Network (NEON), showing Enhanced Vegetation Index (black point), smoothed Enhanced Vegetation Index (green line), period of green-up (green shade), and extracted 50% green-up time (vertical green line), using two trees at Harvard Forest & Quabbin Watershed (HARV, 42.53691° N, 72.17265° W) and Oak Ridge (ORNL, 35.96413° N 84.28259° W) sites as examples.

This empirical method of defining green-up/down time has been widely applied to remote-sensing data in order to be compatible with different plant functional types with various seasonality that exhibit intra-annual changes in greenness (Moon et al., 2021).

We tested the correlation between the 50% green-up time and the flowering time measured by the day of flower onset in the corresponding year from 2018 to 2022. We assessed the Pearson correlation coefficients and the significance of the correlations across all sites.

#### 2.3.2 Inferring pollen phenology on the city level

To infer city-level pollen phenology from tree-level vegetative phenology monitored by PlanetScope, we developed the following nonparametric algorithm with two tuning parameters (Fig. 6, Algorithm 1, Eqn. 3–7). We first extracted the timing of green-up/down events for individual trees at various thresholds based on their EVI curves. These individual events were then upscaled to city-level vegetative phenology. Next, we applied various time lags to vegetative phenology to derive city-level pollen phenology. Finally, we optimized thresholds and lags with air-sampled city-level pollen phenology.

**Figure 6.**
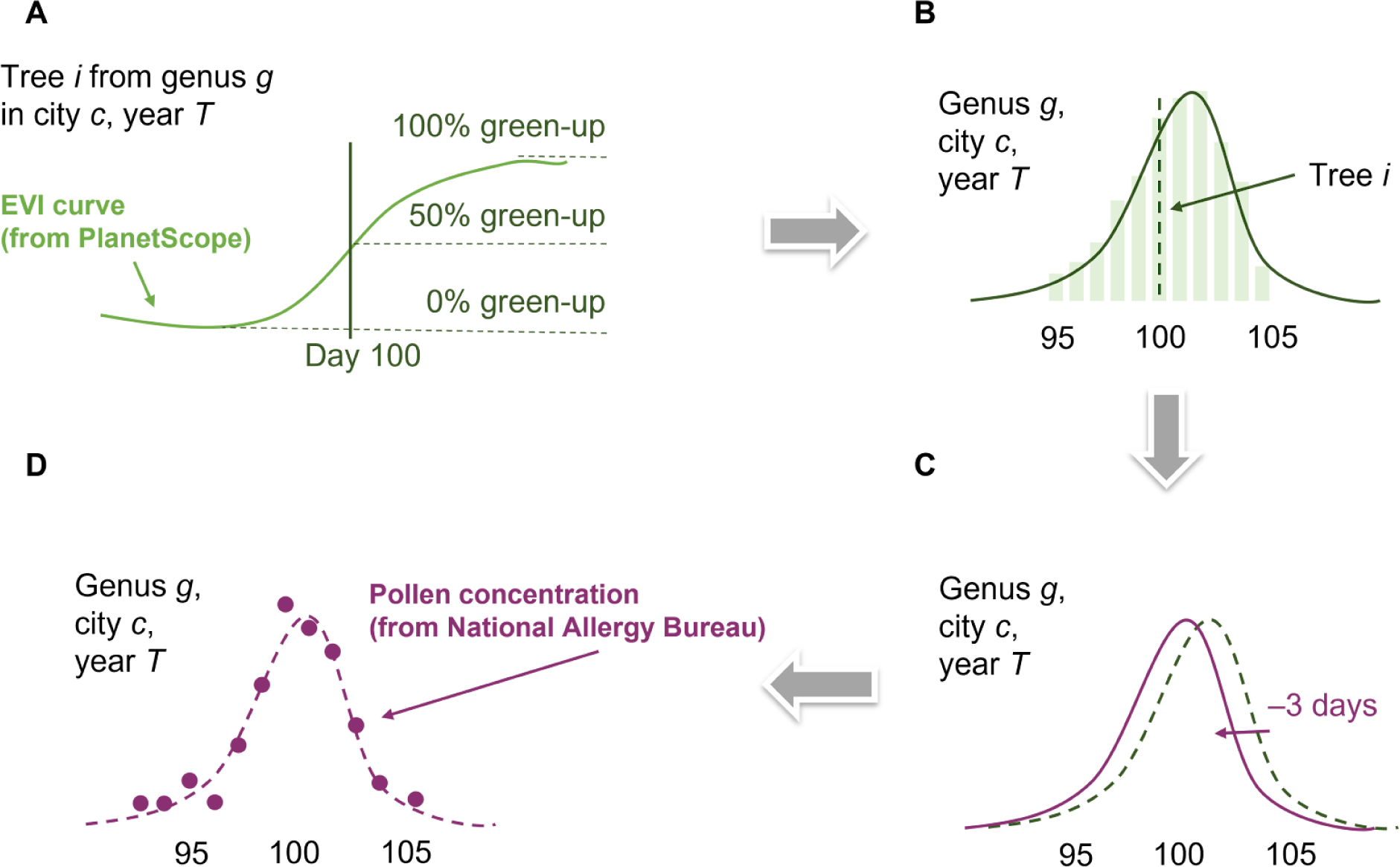
Nonparametric algorithm for inferring pollen phenology from vegetative phenology derived from PlanetScope. The model has four main steps: **(A)** Extract tree-level green-up/down date at threshold *θ_g_* (e.g., *θ_g_* = 50%). **(B)** Upscale to city-level leaf phenology. **(C)** Shift to city-level pollen phenology with leaf-pollen lag *δ_gc_* (e.g., *δ_gc_* = –3 days). **(D)** Compare with city-level pollen concentration.

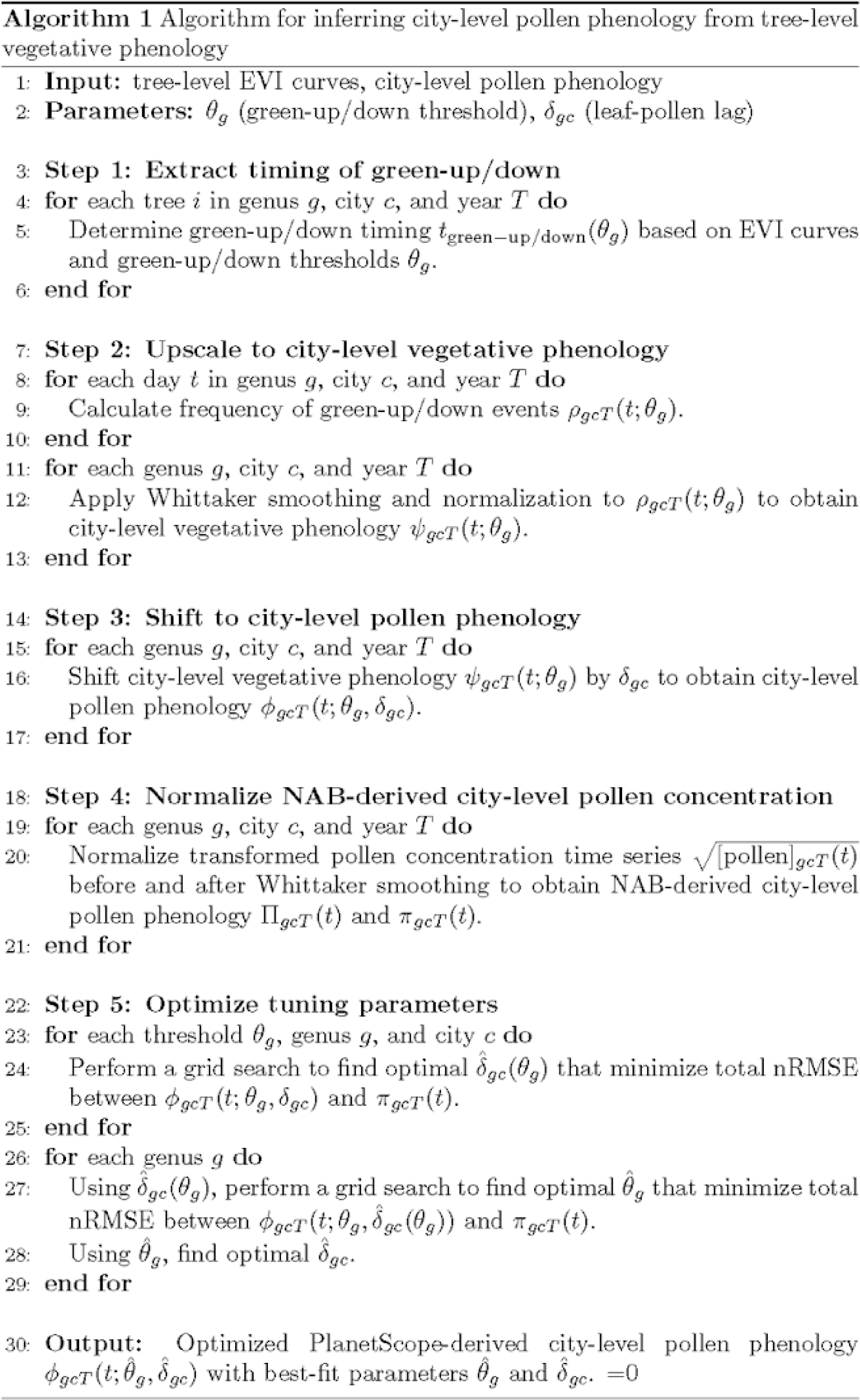

1. We first extracted the timing of green-up/down for all trees of interest as the day of year when their growing season EVI curves first cross the green-up/down threshold *θ_g_* (Eqn. 3). We used a similar algorithm to that described in section 2.3.1 (Fig. 5). The end of a green-up phase was determined as the day of year when EVI reaches the maximum in the growing season. The start of a green-up phase was then determined as the day of year when EVI is at the minimum, prior to the end of the green-up phase. Similarly, the start of a green-down phase was determined as the day of year when EVI reaches the maximum in the growing season; the end of a green-down phase was then determined as the day of year when EVI is at the minimum, after the start of the green-down phase. We then determined the timing of green-up/down at multiple thresholds, including 30%, 40%, 50%, 60%, and 70% green-up for genera that flower in the spring (all except late-flowering *Ulmus* spp.), and 70%,…, 30% green-down for late-flowering *Ulmus* spp.

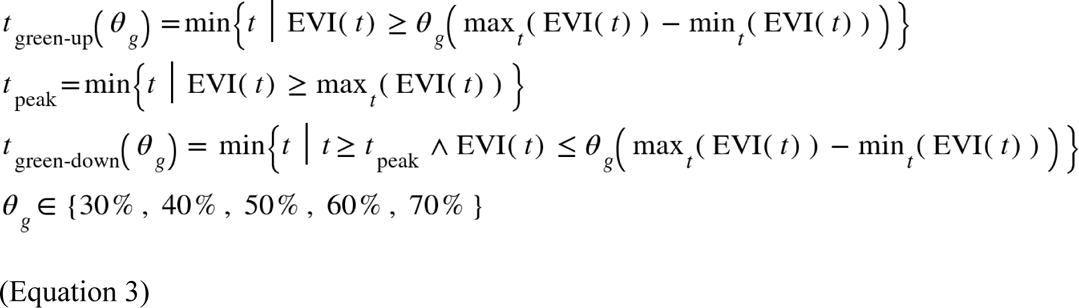

Note that the first tuning parameter in our model is *θ_g_*, the green-up or green-down threshold in the growing season EVI time series that is used to obtain green-up/down time. We allowed the threshold to vary by genus *g*.

1. Given that our pollen concentration data for validation are on the city level rather than tree level, we upscaled the tree-level green-up/down time to the city-level green-up/down frequency 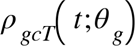 by summarizing the frequency of inferred green-up/down events at green-up/down threshold *θ_g_* on day *t* in a given genus, city, and year. We then applied a weighted Whittaker smoothing and a normalization such that it sums up to one over all days in a year (Eqn. 4).

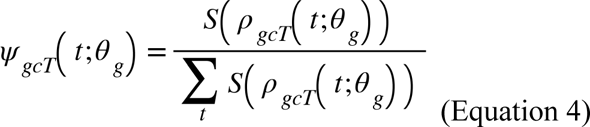

The resulting city-level vegetative phenology 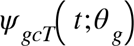 is conceptually similar to a probability density function of observing green-up/down events.

1. Building on the biophysical and empirical relationships between vegetative and reproductive phenology, we assumed that the time of spring pollen emission and the time of leaf-out of a tree has a relatively stable time lag given a specific climate (Buonaiuto & Wolkovich, 2021; Ma et al., 2021; Guo et al., 2023). Late-flowering *Ulmus spp.* is an exception (Wozniak & Steiner, 2017), with little knowledge of the mechanisms of their flowering phenology. We assumed that their flowering time is associated with the senescence phases of vegetative development, similar to other late-flowering species (Rojo et al., 2022).

We shifted the city-level vegetative phenology 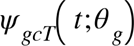 to the city-level pollen phenology 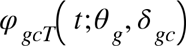 by applying leaf-pollen lags *δ_gc_* (Eqn. 5). We allowed leaf-pollen lags to range from –90 days to 90 days (± 3 months), at the interval of 1 day. We allowed the lag to vary by both genus and city. We acknowledge that this method simplifies the duration of pollen emission to a single pollen emission date (Dahl et al., 2013), hence only capturing the variations in city-level pollen concentration caused by the variations in pollen emission among trees but not within individual trees.

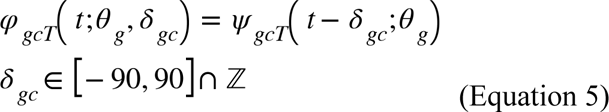

Note that the second tuning parameter in our model is *δ_gc_*, which is the time lag between the timing of leaf phenology and pollen phenology.

1. We scaled the air-sampled city-level pollen concentration to one comparable to PlanetScope-derived city-level pollen phenology, providing city-level pollen phenology for calibration and assessment. We performed normalization to transformed pollen concentration time series 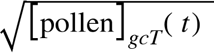 to remove the spatiotemporal differences from pollen sampling methods, local vegetation cover, and pollen productivity. We normalized pollen concentration before and after Whittaker smoothing to obtain NAB-derived city-level pollen phenology 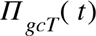 and 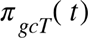 for model assessment and calibration, respectively (Eqn. 6).

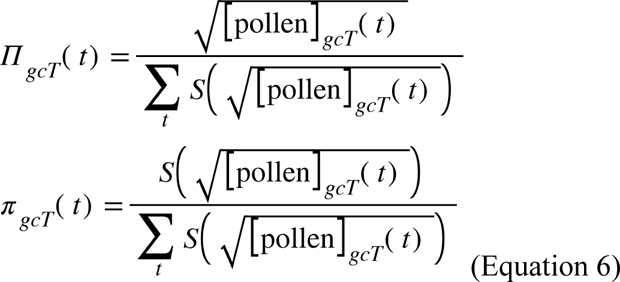

1. We assume that NAB-derived city-level pollen phenology 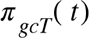 matches PlanetScope-derived city-level pollen phenology 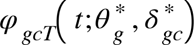with optimized green-up/down threshold 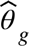 and optimized leaf-pollen lag 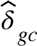. Therefore, we performed a two-step grid search to optimize *θ_g_ and δ_gc_* based on the normalized root mean square error (nRMSE) between PlanetScope and NAB-derived pollen phenology (Eqn. 7). Here, we normalized RMSE by the range of NAB-derived city-level pollen phenology 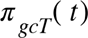 (Jeon et al., 2018; D. Singh & Singh, 2020). We first selected the leaf-pollen lag 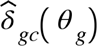 for each green-up/down threshold *θ_g_*that minimized total nRMSE for each genus and city. We then selected the green-up/down threshold 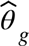 that minimized total nRMSE for each genus. We used 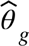 to find 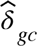.

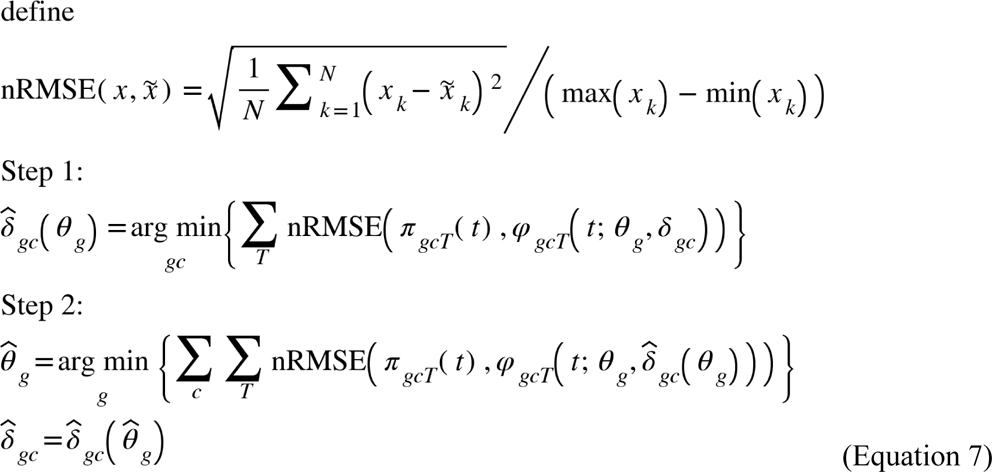

The PlanetScope-derived vegetative phenology modified by the optimized threshold and lags 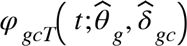 was considered the optimized PlanetScope-derived city-level pollen phenology.

We assessed the accuracy of PlanetScope-derived city-level pollen phenology 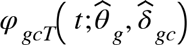 using its Spearman correlation coefficients and nRMSE with NAB-derived city-level pollen phenology 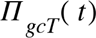. Notably, we compared the Spearman correlation coefficients and RMSE for inferring pollen phenology both in-sample and out-of-sample.

In-sample tests assessed the ability of the PlanetScope method to characterize variations in pollen phenology, whereas out-of-sample tests assessed the effectiveness of the PlanetScope method to infer pollen phenology for cities with no prior pollen concentration observations. For in-sample tests, all cities were used in the optimization of parameters. For out-of-sample tests, we conducted leave-one-out cross-validation. Specifically, we removed a random city from the training dataset at a time and optimized threshold and lags in the remaining cities. To predict the lag for the city held for validation, we assumed a linear relationship between the lag and the climate of the city. To avoid overfitting, we used a simple linear relationship with one predictor representing the long-term temperature of each city. Specifically, we used the mean annual temperature in the TerraClimate Climatologies (1981-2010) dataset (Abatzoglou et al., 2018). With an optimized threshold and a predicted lag, we subsequently inferred pollen phenology from vegetative phenology at the city held for validation. As the out-of-sample tests rely on extrapolation over a climatic gradient, we could only implement them for *Quercus* spp. that were present in all seven cities studied.

#### 2.3.3 Inferring pollen phenology on the tree level

Following the calibration of green-up/down threshold and leaf-pollen lag using NAB-derived city-level pollen phenology, we then scaled back down to infer tree-level pollen phenology using PlanetScope-derived vgreen-up/down time 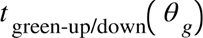 (Eqn. 3). We followed the same assumption used in the previous analysis that the time of peak pollen emission and the time of green-up/down of a tree has a time lag, optimized for each genus and city (Eqn. 8).

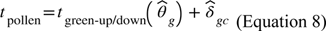

We were able to map the inferred peak pollen emission time of trees in a specific genus, city, and year *t*_pollen_. This allowed us to summarize and visualize intra-urban variations in pollen phenology. Given that NAB airborne concentration data used were collected at one station per city, future validation of PlanetScope-derived pollen allergy landscape with *in-situ* pollen phenology data at finer resolution is warranted.

All data analyses were performed in *R* v. 4.2.0 (R Core Team, 2024).

## 3 Results

### 3.1 PlanetScope-derived vegetative phenology correlates with flowering phenology

We found significant correlations between PlanetScope-derived 50% green-up time and ground-observed flower onset time in wind-pollinated species that were well-sampled (≥ 50 records) across NEON sites in CONUS (Fig. 7). There were significant positive correlations in four out of five species studied, including *Acer pensylvanicum* (striped maple), *Acer rubrum* (red maple), *Quercus montana* (chestnut oak), and *Quercus rubra* (red oak). For the other species, *Quercus stellata* (post oak), which was sampled in only one NEON site, we observed a significant negative correlation. The correlation was limited within sites, where phenological variations are small. These correlations between Planet-derived green-up and ground-observed flower onset were consistent with ground-observed leaf onset and ground-observed flower onset (Fig. S6).

**Figure 7.**
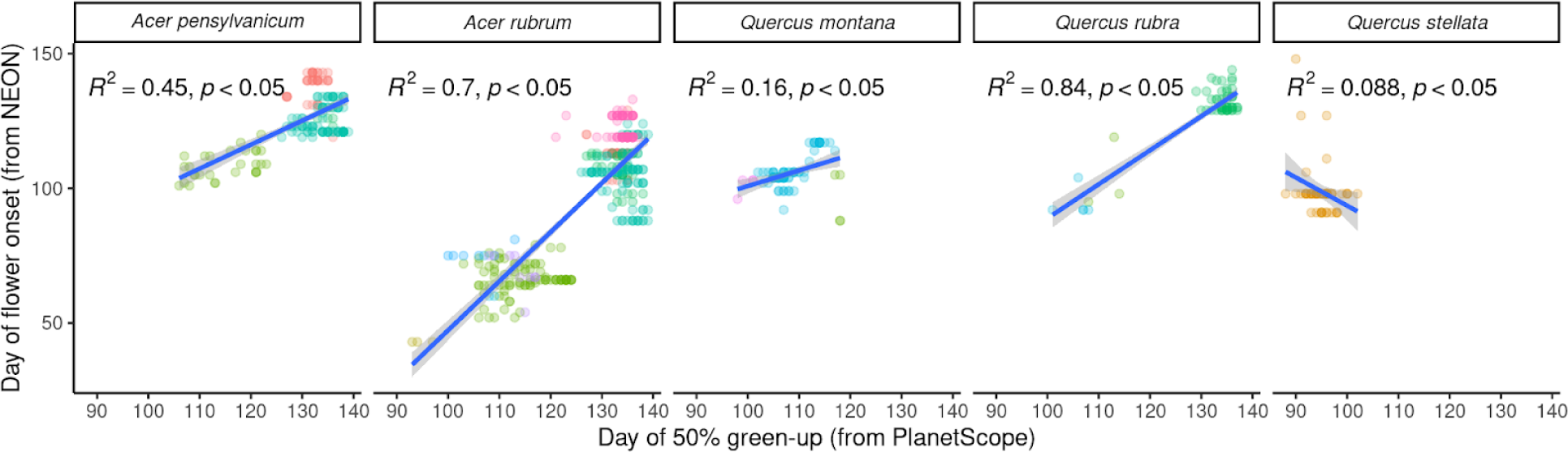
Correlation between 50% green-up time from PlanetScope and flower onset time from the National Ecological Observatory Network (NEON). Different colors indicate NEON sites.

### 3.2 PlanetScope-derived vegetative phenology predicts city-level pollen phenology in-sample and out-of-sample

We were able to infer city-level pollen phenology from PlanetScope-derived tree-level vegetative phenology at a reasonable accuracy with optimized green-up/down thresholds for each genus and further the leaf-pollen lag for each city (Fig. 8, Figs. S7, S8, S9, S10). Among all 195 combinations of genus, city, and year, the in-sample accuracy of our PlanetScope method achieved a Spearman correlation of 0.614 (median, 95% interval: 0.133–0.861) and a nRMSE of 14.6% (9.07%–40.7%). In in-sample tests, 178 out of 187 (95.2%) Spearman correlations were statistically significant (*p* ≤ 0.05). Across genera, the highest and lowest Spearman correlations and nRMSE were seen in *Quercus* spp. (correlation: 0.751, 0.491–0.918; nRMSE: 13.5%, 9.67%–32.1%) and in *Acer* spp. (correlation: 0.514, 0.131–0.736; nRMSE: 18.9%, 10.0% –40.8%).

**Figure 8.**
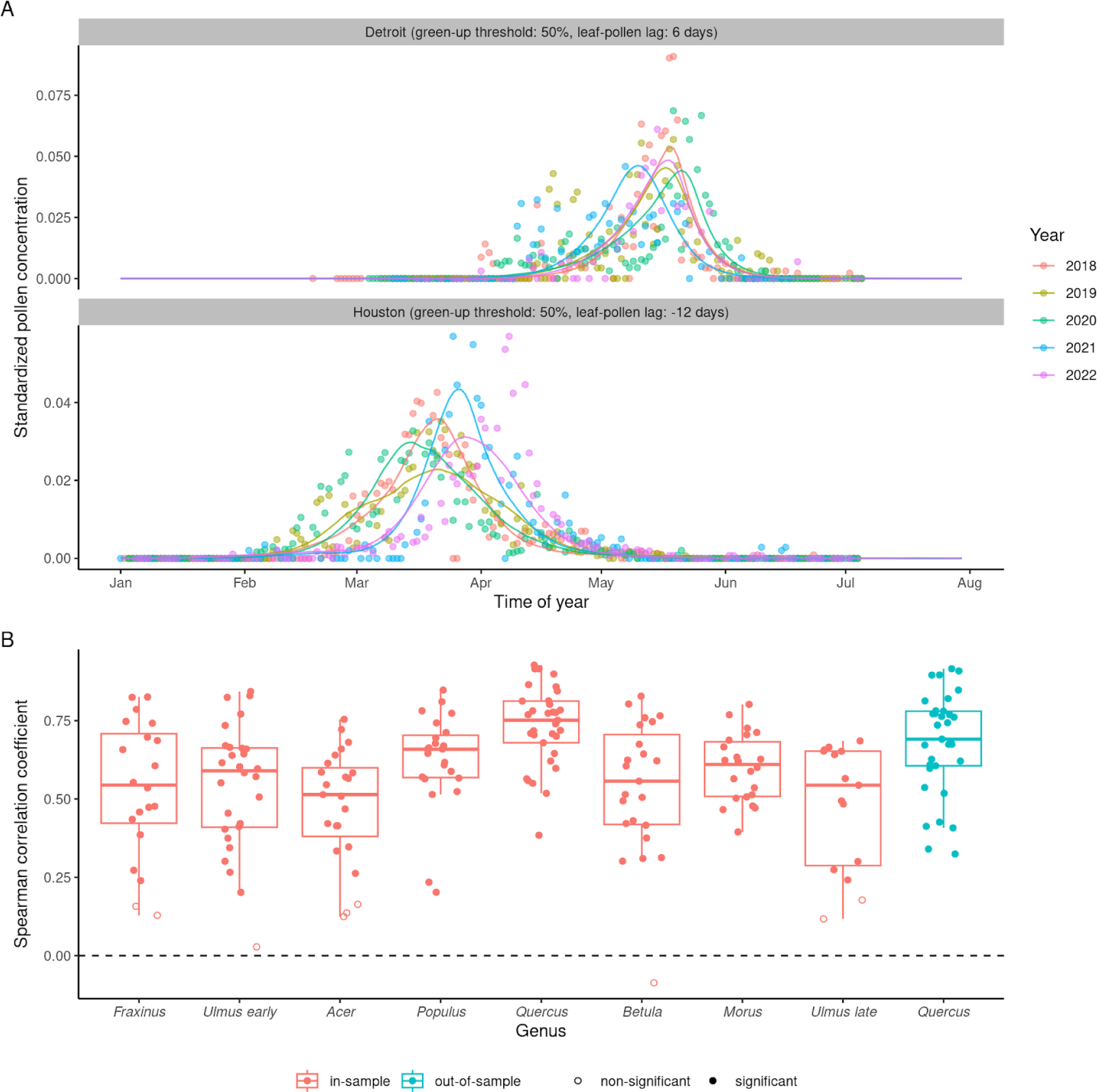
Comparing city-level pollen phenology derived from PlanetScope and from airborne pollen concentration monitored at the National Allergy Bureau pollen monitoring stations. **(A)** Pollen phenology inferred from PlanetScope-derived vegetative phenology tuned to the optimal green-up/down thresholds and leaf-flower lags (lines) compared to pollen phenology inferred from airborne pollen concentration (points). Pollen phenology from both data sources was standardized to probability density distribution within each city and year to allow comparison. Here we show examples of *Quercus* spp. (oak) pollen phenology in two cities in the south (Houston) and north (Detroit) of CONUS. **(B)** Accuracy of inferring pollen phenology with the PlanetScope method, both in-sample (fitting model with data from all cities) and out-of-sample (leave-one-out cross-validation), measured by Spearman correlation, indicating the level of significance (*p* ≤ 0.05).

To test if our method can be generalized to locations without prior pollen concentration data, we performed an out-of-sample test for *Quercus* spp. that was present at seven cities, assuming a linear relationship between leaf-pollen lag and long-term climate of the city (Fig. S11). Our PlanetScope method achieved an out-of-sample Spearman correlation of 0.691 (0.337–0.910), with all 33 correlations being statistically significant, and an out-of-sample nRMSE of 14.5% (9.82%–36.0%). Out-of-sample performances were comparable to in-sample performances.

### 3.3 PlanetScope-derived vegetative phenology informs within-city variations in pollen phenology

Beyond the promise of extrapolating over a large spatial scale, we also explored the potential of leveraging the fine spatial resolution of PlanetScope time series to map the spatial details of pollen concentration within cities. We found that both the distributions of spring leaf green-up time and pollen emission time among trees of interest in a city differ from often-assumed Gaussian kernels, characterized by asymmetric peaks and sometimes multiple peaks (Fig. 9A). Because of the different relationships between vegetative and reproductive phenology among genus, we inferred that the distributions of pollen emission time among trees of interest were different from the distributions of spring leaf green-up time, further altering the shape of pollen peaks (Fig. 9A). We mapped the inferred pollen emission among trees, showing the pollen allergy landscape with spatial variations within cities (Fig. 9B).

**Figure 9.**
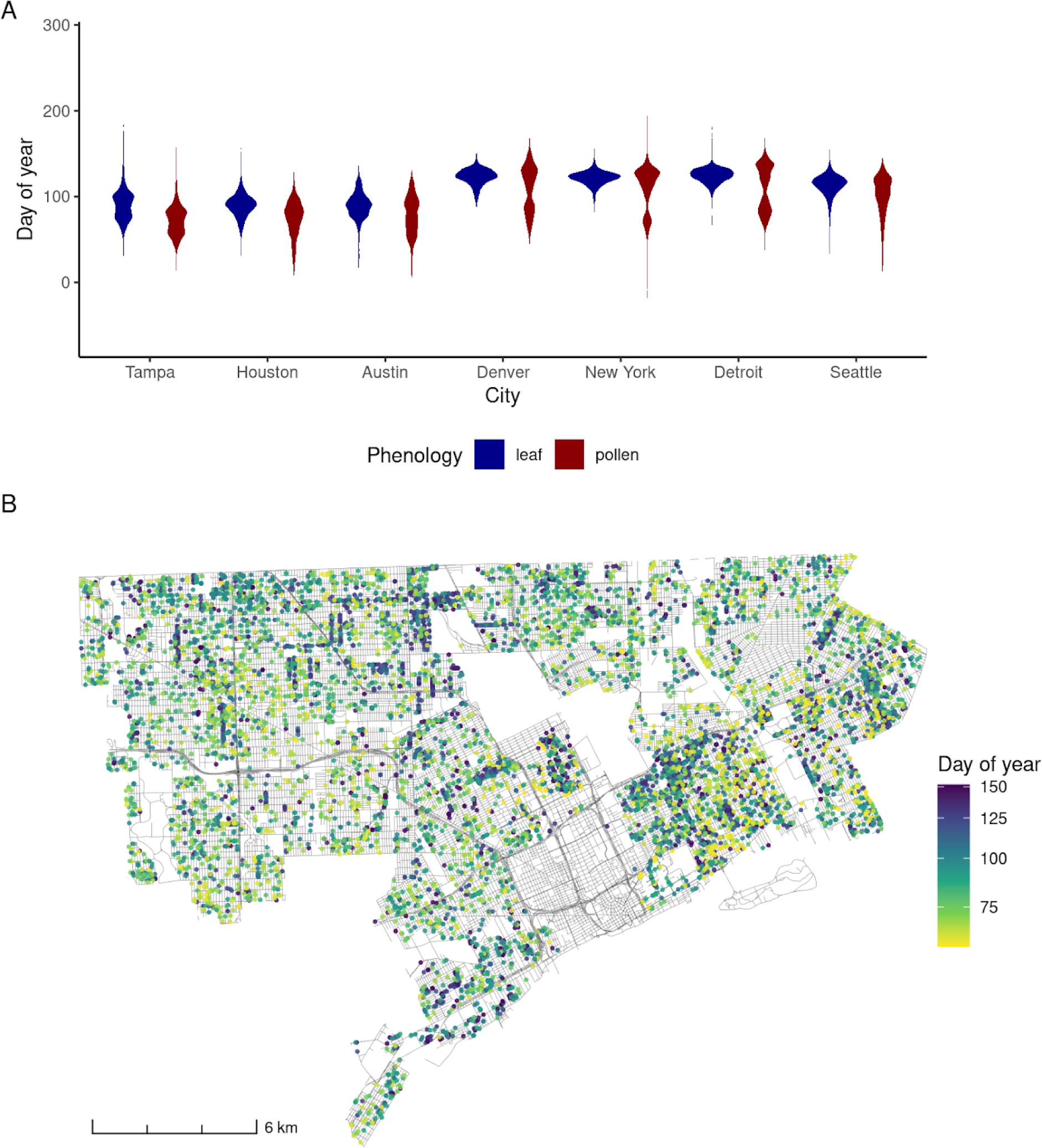
**(A)** Distributions of PlanetScope-derived tree-level spring leaf green-up time and peak pollen emission time within cities over all years. Cities are ordered in latitudes, from low to high. **(B)** Map of PlanetScope-derived peak pollen emission time in Detroit street trees in the spring of 2018.

## 4 Discussion

In this study, we developed a workflow to infer flowering and pollen phenology from PlanetScope-derived vegetative phenology, validated by *in-situ* phenological observations and measurements of airborne pollen concentrations. On the tree level, PlanetScope-derived green-up time was correlated with flower onset time. On the city level, PlanetScope-derived green-up/down time at an optimized threshold and shifted by a time lag can be used to characterize pollen phenology, with the possibility to predict out-of-sample in cities without pollen concentration observations. Further, we demonstrated the power of PlanetScope time series in mapping the pollen allergy landscape within cities. This study reveals the potential of high spatiotemporal resolution remote sensing data for modeling the reproductive phenology of wind-pollinated plants and mapping allergenic pollen exposures on large scales and in great spatial details.

### 4.1 Relationship between vegetative and reproductive phenology

In both tree- and city-level analyses, we showed the link between vegetative and reproductive phenology, either flower onset or pollen emission. Similar relationships have been widely supported in previous studies but on large spatial and taxonomic scales. Across locations within the geographical range of species distribution, remotely sensed land surface phenology explained variations in flowering and pollen season onset (Høgda et al., 2002; Karlsen et al., 2008). The regional-level leaf-flower correlation can be explained mechanistically, that spring flowering and green-up times are both functions of accumulated temperature for many species (Cook et al., 2012). Across species, there was a significant positive correlation between the timing of the first flower and the timing of the first leaf, after controlling for phylogeny (Davies et al., 2013; Du et al., 2017). The phylogenetic conservatism of this leaf-flower correlation suggests deep evolutionary linkages between phenological responses to a common set of environmental cues (Davies et al., 2013; Delbart et al., 2015; Du et al., 2017).

The optimal timing of plant life-history events in temperate ecosystems has been predicted to be constrained by the trade-off between harsh abiotic conditions and high biotic competition (Iwasa & Levin, 1995; Post, 2013; CaraDonna & Bain, 2016). This theory applies to both vegetative and reproductive phenology. The spring leaf-out of most temperate woody plant species is a result of balancing the advantages of a longer growing season with the risks from frost damage, mechanistically controlled by a suite of winter temperature, spring temperature, and photoperiod cues (Polgar & Primack, 2011). Flowers, being more sensitive than leaves to frost damage (CaraDonna & Bain, 2016), are also highly controlled by similar environmental cues (Wang et al., 2020). In addition, the vegetative and reproductive phenology of wind-pollinated species might also be directly constraining each other, as they often flower before the full closure of the canopy for efficient pollen dispersion (Buonaiuto et al., 2021; Buonaiuto & Wolkovich, 2021).

Despite established leaf-flower correlation across space and species, this relationship has rarely been examined on the individual tree level (but see Primack, 1985). Here, we showed that individual trees that green up earlier also tend to flower early, which may be attributed to the shared response of both phenology to spatial variations in microclimates (Katz et al., 2019), or genetic differences in plant development among trees (Primack, 1985). By revealing a leaf-flower correlation that holds on a scale smaller than previously known, we suggest the presence of fine-scale mechanisms for this correlation in addition to climatic and phylogenetic control. Such insight into the relationship between phenological events might inform the integration of leaf phenology in the mechanistic model for flower and pollen phenology.

### 4.2 Detecting tree-level phenology with PlanetScope

Validated by *in-situ* phenological observations, we closely examined the often-assumed potential of PlanetScope to detect tree-level phenology (Cheng et al., 2020; Wu et al., 2021; Zhao et al., 2022). Across NEON sites, PlanetScope-derived phenometrics captured tree-level variations in the onset of maple and oak flowers. This result is consistent with the finding that PlanetScope-derived phenometrics explained tree-level variations in the leaf onset time of deciduous trees across NEON sites (Zhao et al., 2022) and in leaf senescence time at a PhenoCam site in a temperate forest (Wu et al., 2021).

Nevertheless, we noted that tree-level correlations were contributed mainly by cross-site rather than within-site correlations (Fig. 7). The weak within-site correlations were also observed in Zhao et al. (2022) and were explained with both the uncertainty of extracting phenometrics from PlanetScope time series and inconsistencies in field phenological observations. An additional reason is that the correlations between leaf onset time and flower onset time, a premise of our method, might be weaker within the site compared to across sites (Fig. S6).

The ability to detect tree-level phenology using PlanetScope time series is highly relevant from both a scientific and a practical standpoint. On one hand, there are large variations between the phenology of trees within a population, often with a conserved order of phenological events among trees repeating from year to year, referred to as “phenological rank” (Delpierre et al., 2017). Such tree-level phenological variations could arise from multiple factors, including phenological variations in phenotype, microclimate, and local resource availability (Vitasse et al., 2021). PlanetScope-derived phenology will allow a better understanding of ecological factors that underlie phenology. In practice, tree-level variations in pollen phenology contribute to spatial heterogeneity of pollen concentration within a city (Katz et al., 2019).

While current pollen concentration data and pollen models are largely confined to regional or city scales, PlanetScope-derived pollen phenology has the potential to empower process-guided pollen models with a higher spatial resolution. Here, we demonstrate how our PlanetScope-derived green-up time tuned to the optimal thresholds and lags, integrated with street-tree inventory, allowed us to map the spatial heterogeneity of PlanetScope-derived pollen emission time in a given city and time (Fig. 9). Such maps showing intra-urban phenological variations can inform decision-making such as time and location of outdoor activities. Future validation with spatiotemporal pollen phenology data on sub-city scales is warranted (Zapata-Marin et al., 2022).

The method we employed to extract tree-level phenology here using coordinates of censused trees is highly scalable, but several improvements can be made when working on smaller scales, or if automated for scaling up. Manual segmentation of tree crowns from drone imagery has been used to delineate polygons to extract phenological signals from PlanetScope (Wu et al., 2021; Zhao et al., 2022). Such tree crown segmentation could be automated in the presence of co-registered airborne RGB, LiDAR hyperspectral imagery, and high-quality training data (Weinstein et al., 2021). Given uncertainties in effective footprint size and geolocation accuracy of PlanetScope pixels, small trees (e.g., with a canopy diameter < 3 m) might not be suitable to be extracted for the phenological signal from PlanetScope. The size of trees could be controlled if there exists information on tree diameter, height, or age in tree inventories; alternatively, this could be achieved by segmentation of tree canopy.

### 4.3 Using PlanetScope time series to inform pollen phenology

In the city-level analysis using pollen concentration data, we proposed a novel workflow to predict pollen phenology from PlanetScope time series. We compared the performance of our method to previous machine learning models that do not explicitly account for phenology and a study that directly accounts for flowering phenology. On the one hand, our in-sample nRMSE of 14.6% and out-of-sample nRMSE of 14.5% (for *Quercus* spp. only) were highly competitive with machine learning models with nRMSE generally above 20% (Seo et al., 2019; Makra et al., 2023), even though we did not include any environmental predictors to account for climate or weather changes. On the other hand, our performance was comparable to or better than that of the System for Integrated modeLling of Atmospheric coMposition (SILAM), a widely-used process-based modeling system in Europe, which gives an nRMSE of approximately 20% in pollen prediction (Sofiev et al., 2015). Further, we outperformed a previous study that predicted the same NAB pollen concentration dataset with *in-situ* flower observations collected by community scientist volunteers (Crimmins et al., 2023). We achieved higher and more statistically significant Spearman correlations in this study: 58% and 95.2% of Spearman correlations were statistically significant in Crimmins et al. (2023) and this study, respectively; the median Spearman correlation for *Quercus* spp. was 0.49 and 0.751 in Crimmins et al. (2023) and this study, respectively. The competitive performance of our model demonstrated the power of phenology data in pollen models and the great potential of PlanetScope-derived phenology data.

The accuracy of our method arises from the ability of PlanetScope-derived pollen phenology to explain intra-annual and inter-annual variations in pollen concentration. By empirically inferring the continuous change in pollen phenology in a year from the distribution of green-up/down days among trees (Fig. 8, Fig. S11), we moved beyond traditional pollen phenology modeling that relies on annual phenometrics (Clark et al., 2014). Even without explicitly accounting for inter-annual variations in temperature, the PlanetScope method was able to explain some inter-annual variations in pollen phenology with the different times and distributions of green-up/down dates (Fig. 8, Fig. S8). This finding has previously been shown on a coarser resolution, with a remotely-sensed green-up date explaining inter-annual variations in ground-observed flowering dates (Delbart et al., 2015).

Our model was a simplified one, overlooking details such as differential responses of vegetative and reproductive phenology to the environment (Geng et al., 2022) and non-phenological factors that influence pollen concentration, such as pollen dispersion (Latorre, 1999; Robichaud & Comtois, 2021; Zhu et al., 2024). Nevertheless, this model paves the way for building process-guided pollen models with tree-level vegetative and reproductive phenology as key processes (Scheifinger et al., 2013; Katz, Vogt, et al., 2023).

The out-of-sample accuracy of PlanetScope-derived pollen phenology of oak trees (Fig. 8, Fig. S10) suggested the possibility of predicting pollen concentration even at locations with limited prior observations. We suggest four enhancements before operationalizing the proposed workflow to inform decision-making in public health. First, in addition to the tree-level validation of the flowering phenology, we presented, more fine-scale ground truths for pollen phenology are needed (Katz & Batterman, 2020). Examples are newly initiated community science data collection of pollen phenology (Katz, Vogt, et al., 2023) and within-city pollen concentration data (Weinberger et al., 2018). Second, our study was partly limited by the number of cities with publicly available street inventories. To ensure a sufficient number of cities for extrapolation, we were not able to perform out-of-sample tests with genera other than *Quercus spp.* Further model tuning and validation will benefit from obtaining street tree inventories from more cities. Apart from direct requests from cities, operationalized identification of trees with remote sensing data (Morueta-Holme et al., 2024), such as the Auto Arborist dataset (Beery et al., 2022), may be particularly efficient. Third, by focusing on phenology, our method addressed the relative change of pollen concentration but not the absolute magnitude of pollen peaks, which can be achieved by accounting for the total pollen production per tree and abundance and distribution of the taxa of interest (Katz & Carey, 2014; Katz et al., 2020; Zhang & Steiner, 2022). Last, although this study retrospectively demonstrates the inference of pollen phenology, this method can be combined with accurate forecasting of vegetative phenology (Song et al., 2023; Taylor & White, 2020) to achieve near-term predictions for the early warning of pollen season. With around 60 pollen monitoring stations around the US but far more widespread public health concerns on pollen allergy, this study provides the possibility of more spatially equitable access to pollen level forecasting.

## 5 Conclusions

We showed that high spatiotemporal resolution remote sensing data from PlanetScope is highly promising in inferring flowering and pollen phenology from vegetative phenology. On the tree level, PlanetScope-derived green-up time had a significant positive correlation with the field-observed flower onset time of four out of five wind-pollinated species in a large observatory network. We upscaled tree-level PlanetScope-derived vegetative phenology to the city level and accurately inferred pollen phenology, calibrated by a continental-scale high-quality dataset for airborne pollen concentration. This empirical method of inference achieved a median Spearman correlation of 0.606 and a nRMSE of 14.8% across seven wind-pollinated genera and seven cities of interest. For *Quercus* spp. (oak) that was present in all seven cities; our model achieved high out-of-sample accuracy (Spearman correlation: 0.691, nRMSE: 14.5%) comparable with in-sample accuracy (Spearman correlation: 0.751, nRMSE: 13.5%). Our proposed method is promising to describe and predict the pollen phenology of deciduous wind-pollinated tree taxa at locations without prior pollen concentration data. We further demonstrate how PlanetScope-derived vegetative phenology can be used to map variations in pollen phenology within cities (a pollen allergy landscape), calling for future validation with fine-grained *in-situ* data. Using PlanetScope time series, our findings pave the way for the development of generalizable and refined process-based pollen models to inform decision-making in public health.

## CRediT authorship contribution statement

**Yiluan Song:** Conceptualization, Methodology, Data curation, Formal analysis, Writing – original draft, Writing – review & editing. **Daniel S.W. Katz:** Conceptualization, Data curation, Writing – review & editing. **Zhe Zhu:** Conceptualization, Methodology, Writing – review & editing. **Claudie Beaulieu:** Methodology, Writing – review & editing. **Kai Zhu:** Supervision, Resources, Conceptualization, Writing – review & editing.

## Declaration of competing interest

The authors declare that they have no known competing financial interests or personal relationships that could have appeared to influence the work reported in this paper.

## Acknowledgments

The authors would like to thank Drs. Yang Chen and Inés Ibáñez, and members of the Zhu Lab at the University of Michigan, for comments on the manuscript. Yiluan Song was supported by the UCSC Hammett Fellowship, the Microsoft AI for Earth Grant, and the Eric and Wendy Schmidt AI in Science Postdoctoral Fellowship, a Schmidt Sciences program. Kai Zhu and Yiluan Song were supported by the National Science Foundation [grant numbers 2306198 (CAREER)] and Michigan Institute for Data and AI in Society.

## Data availability

We retrieved ground-observed plant phenology data (DP1.10055.001) from the National Ecological Observatory Network (NEON) integrated into the USA National Phenology Network (USA-NPN) using R package *rnpn* on Feb 1, 2024. We obtained pollen concentration data from pollen monitoring stations associated with the National Allergy Bureau. Pollen concentration data were received on Apr 25, 2023 and will not be released due to confidentiality reasons. We downloaded PlanetScope imagery for areas of interest through the Planet API, adapting an R package *planetR*, during Feb 1, 2024 – Mar 14, 2024. Processed data and novel R code will be available upon request.

## Notes

### Competing Interest Statement

The authors have declared no competing interest.

