## Supplementary Information for "Predicting reproductive phenology of wind-pollinated trees via PlanetScope time series"

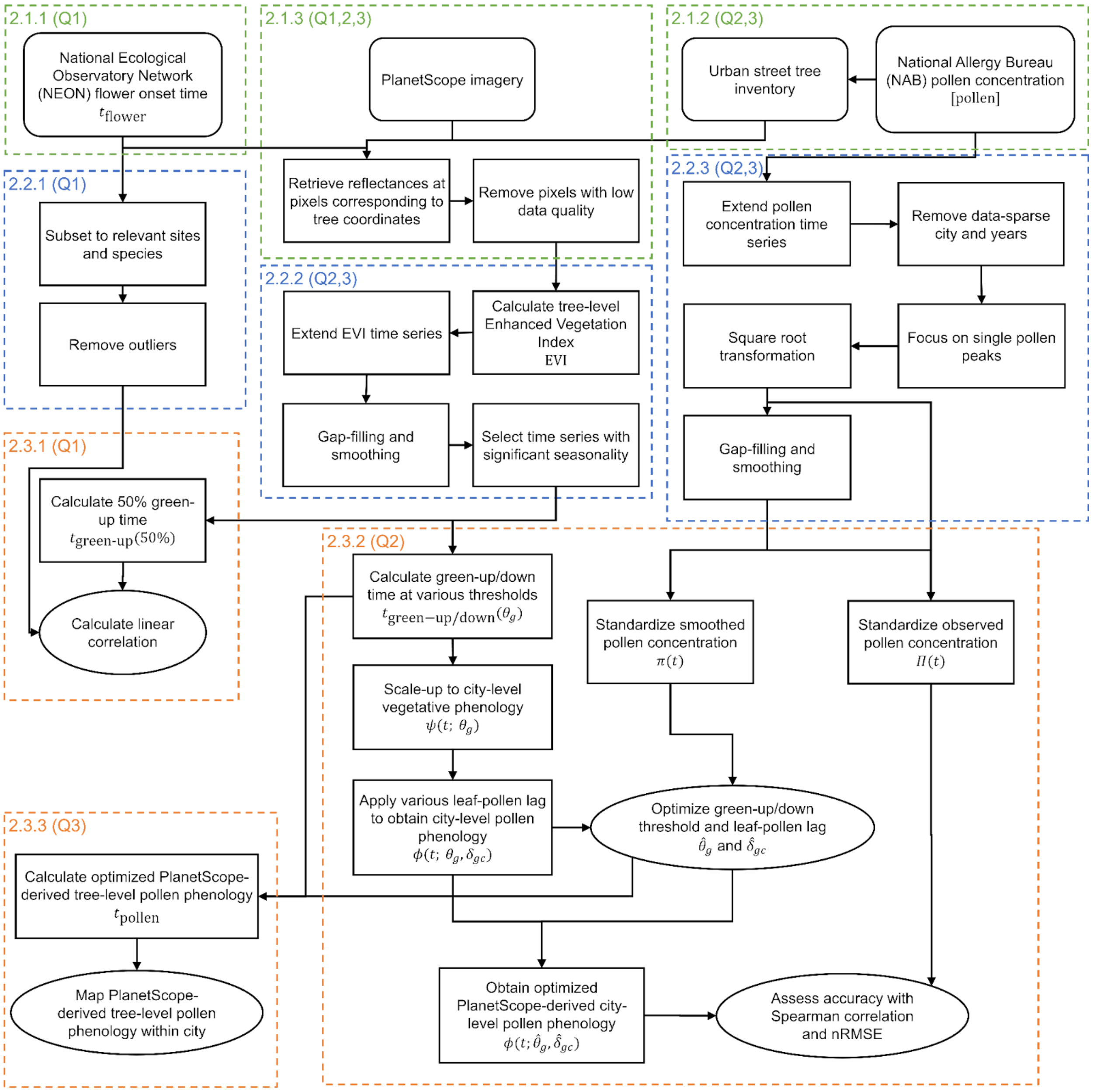


**Figure S1.** Flow diagram of data and methods used in this study, and corresponding research questions. Here, $t$ represents day of year. Indices $g$ and $c$ represent genus and city, respectively. Variable names are explained in equations. Numbers in dashed squares indicate corresponding sections in the main text.

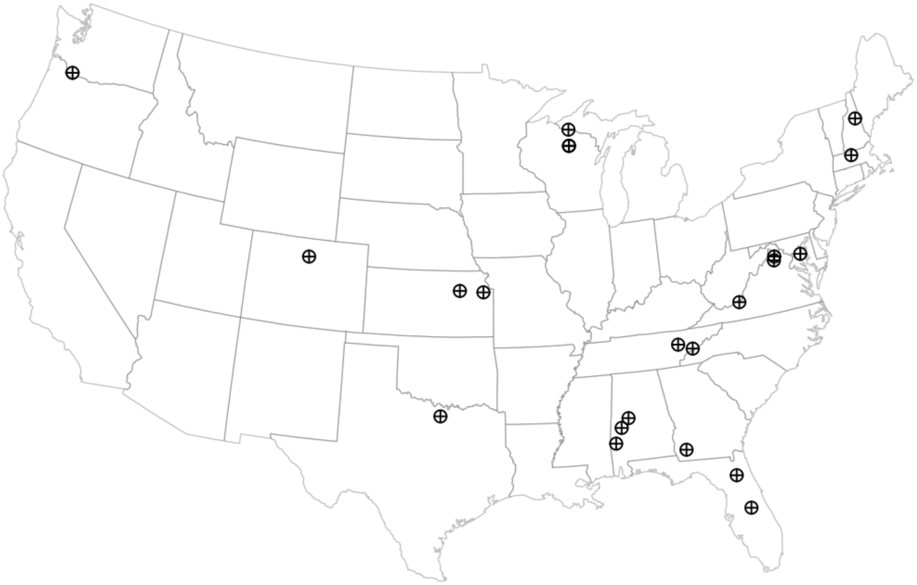

**Figure S2.** Map of National Ecological Observatory Network (NEON) sites used to examine the relationship between PlanetScope-derived leafing phenology and ground-observed flowering phenology.

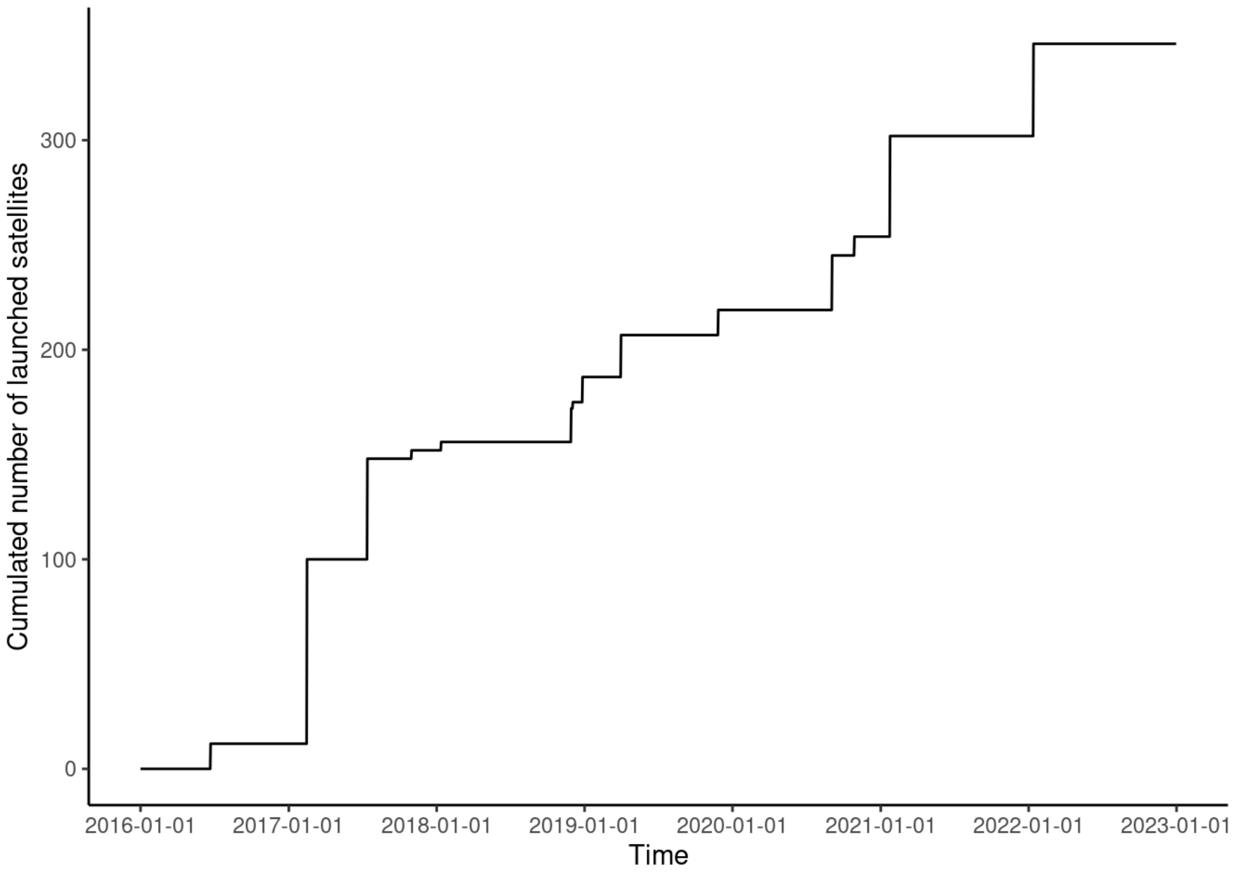


**Figure S3.** Timeline of PlanetScope DOVE satellite launching, with data retrieved from https://earth.esa.int/eogateway/missions/planetscope/description.

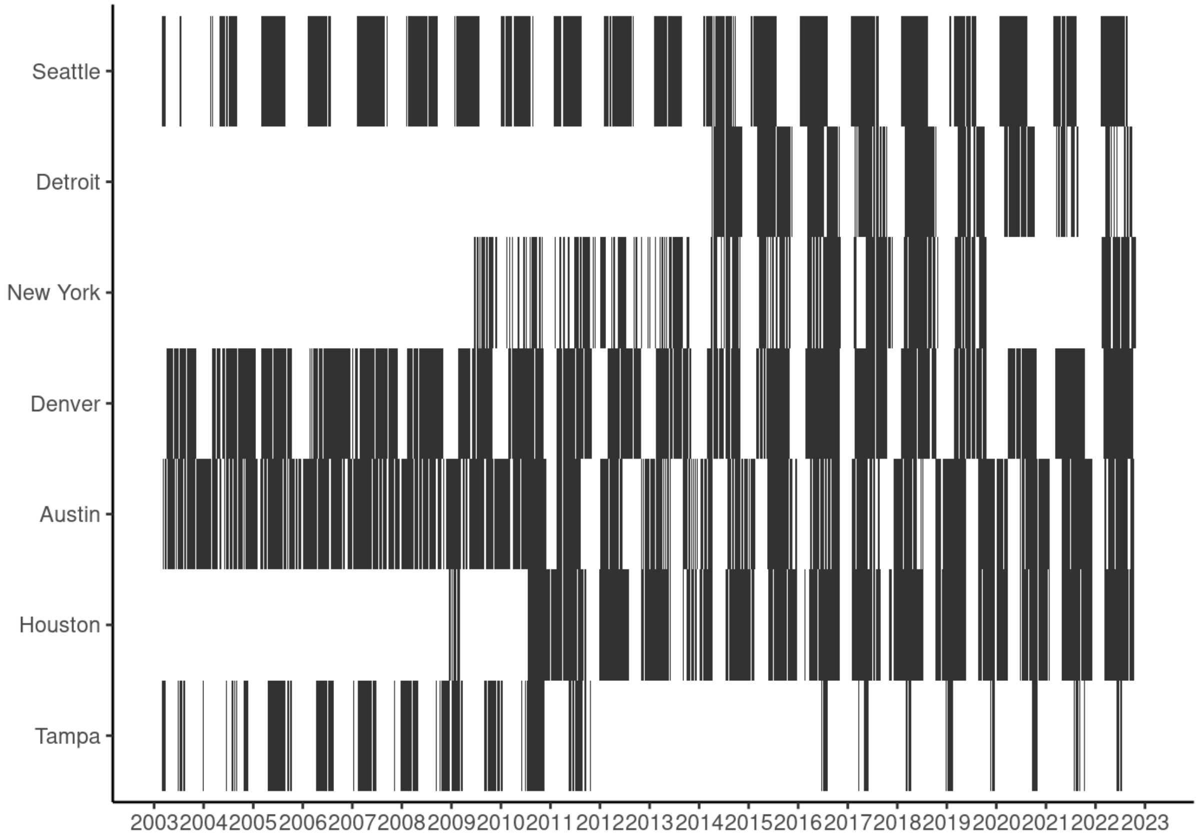


**Figure S4.** Availability of pollen concentration data at study sites, with black color indicating available data. Data were obtained from pollen counting stations associated with the National Allergy Bureau (NAB) on Apr 25, 2023.

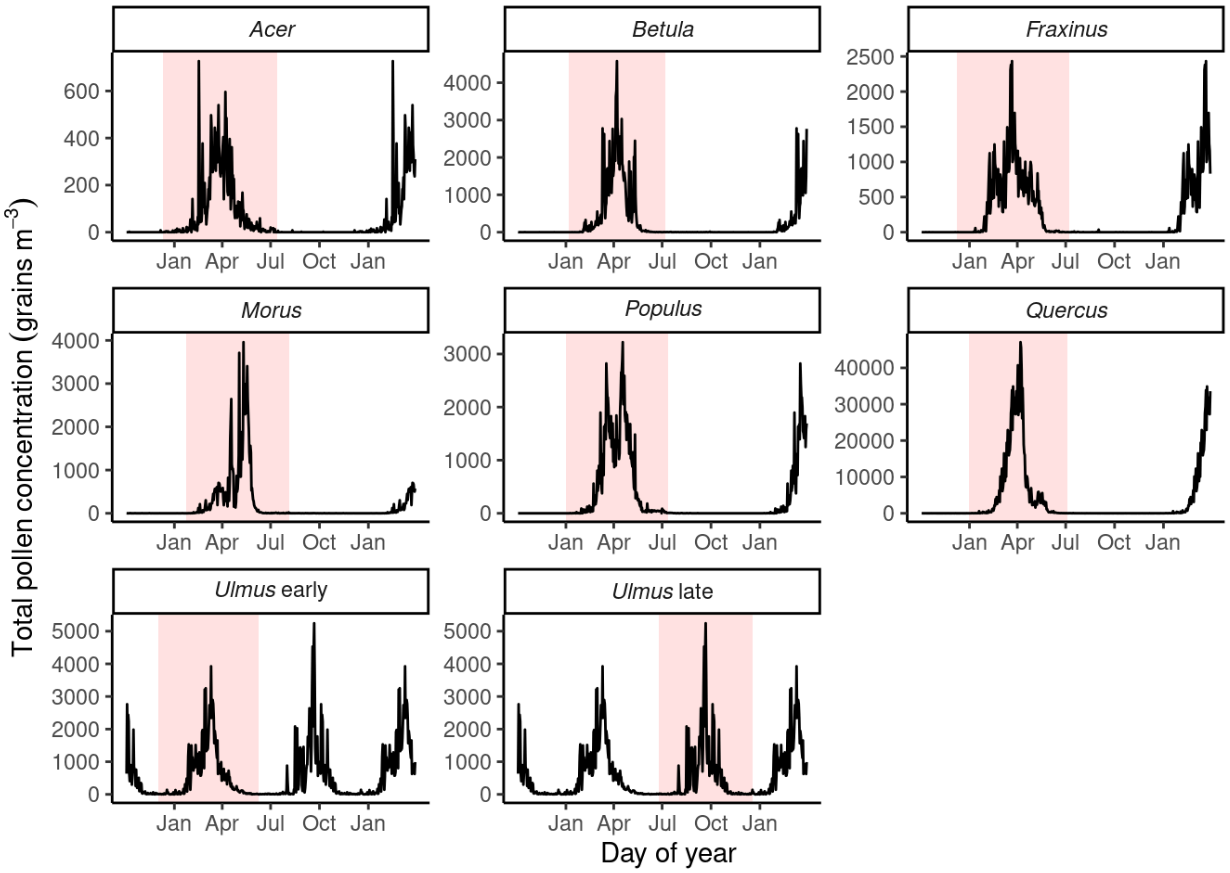


**Figure S5.** Taxa-specific pollen window determined by historical pollen concentration data. The total pollen concentration over all sites and years (black lines) were fitted with Gaussian distributions (red lines). Mean and standard deviations of the Gaussian distributions were used to determine pollen windows (orange rectangle).

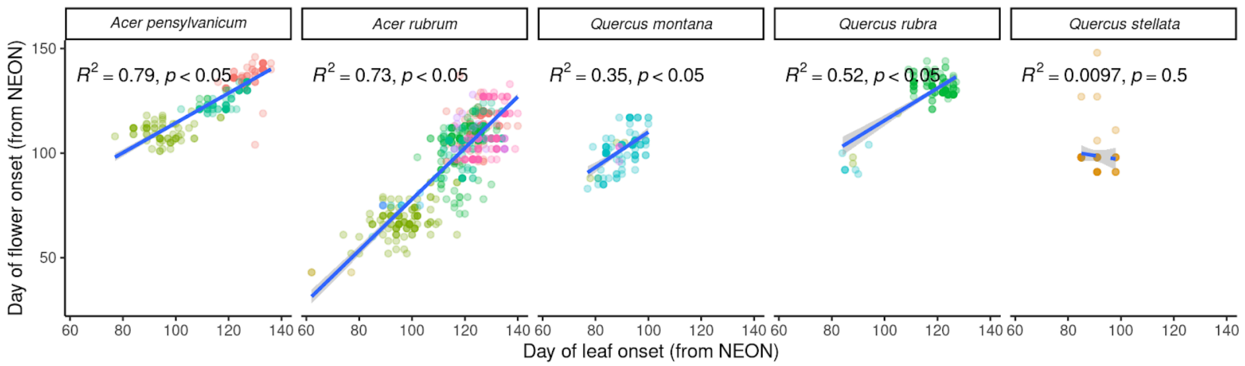


**Figure S6.** Correlation between leaf onset time and flower onset time from the National Ecological Observatory Network (NEON). Different colors indicate NEON sites.

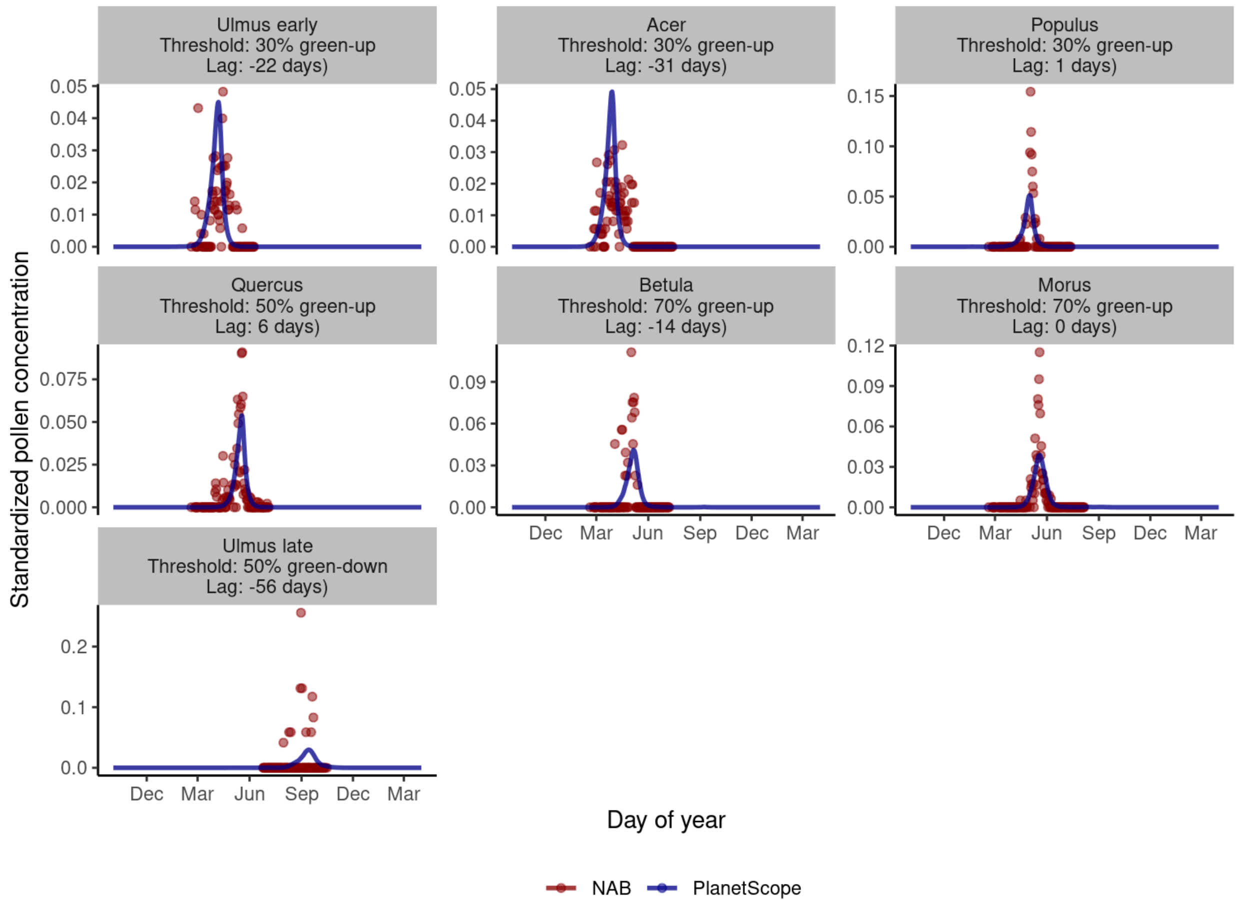


**Figure S7.** Comparing pollen phenology derived from airborne pollen concentration in the National Allergy Bureau (NAB) dataset and pollen phenology derived from PlanetScope time series across seven wind-pollinated genera in Detroit in the year of 2018. Optimized green-up/down thresholds and leaf-pollen lags are shown for each genus.

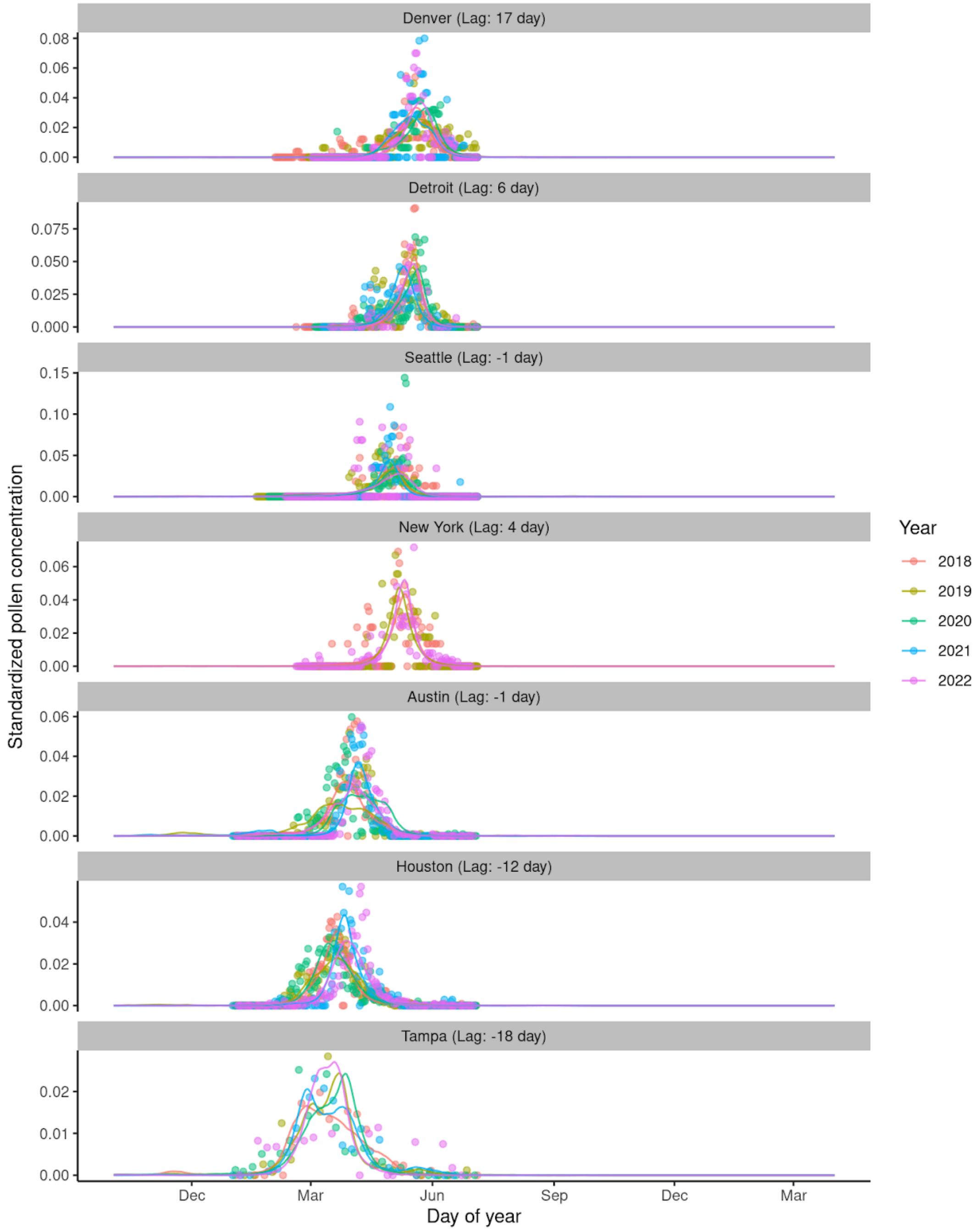


**Figure S8.** Comparing pollen phenology derived from airborne pollen concentration in the National Allergy Bureau (NAB) dataset and pollen phenology derived from PlanetScope time series for *Quercus* spp. across seven cities and five years studied. The green-up threshold was optimized to be 50% and different leaf-pollen lags are shown for each city.

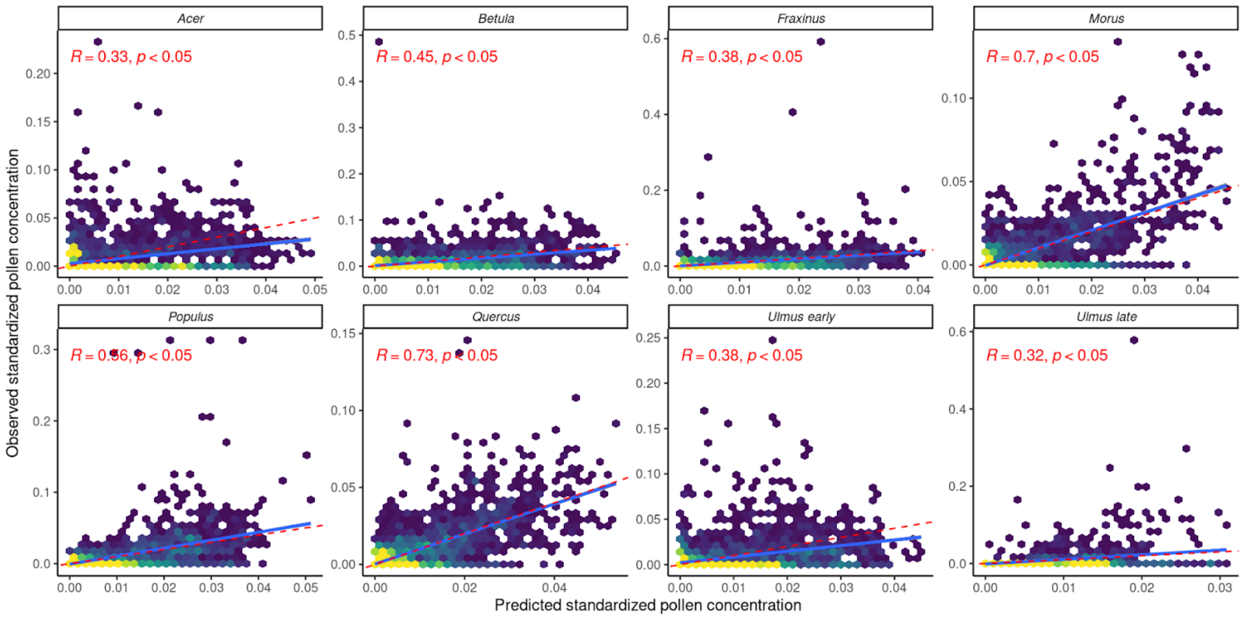


**Figure S9.** Heatmaps showing the correlation between pollen concentration predicted from PlanetScope-derived vegetative phenology and observed at National Allergy Bureau (NAB) pollen counting stations, both after standardization, in all combinations of genera, city, and years. Blue solid lines show linear regression lines and red dashed lines show 1:1 lines. Pearson correlation coefficient (*R*) and level of significance (*t*-test) are shown.

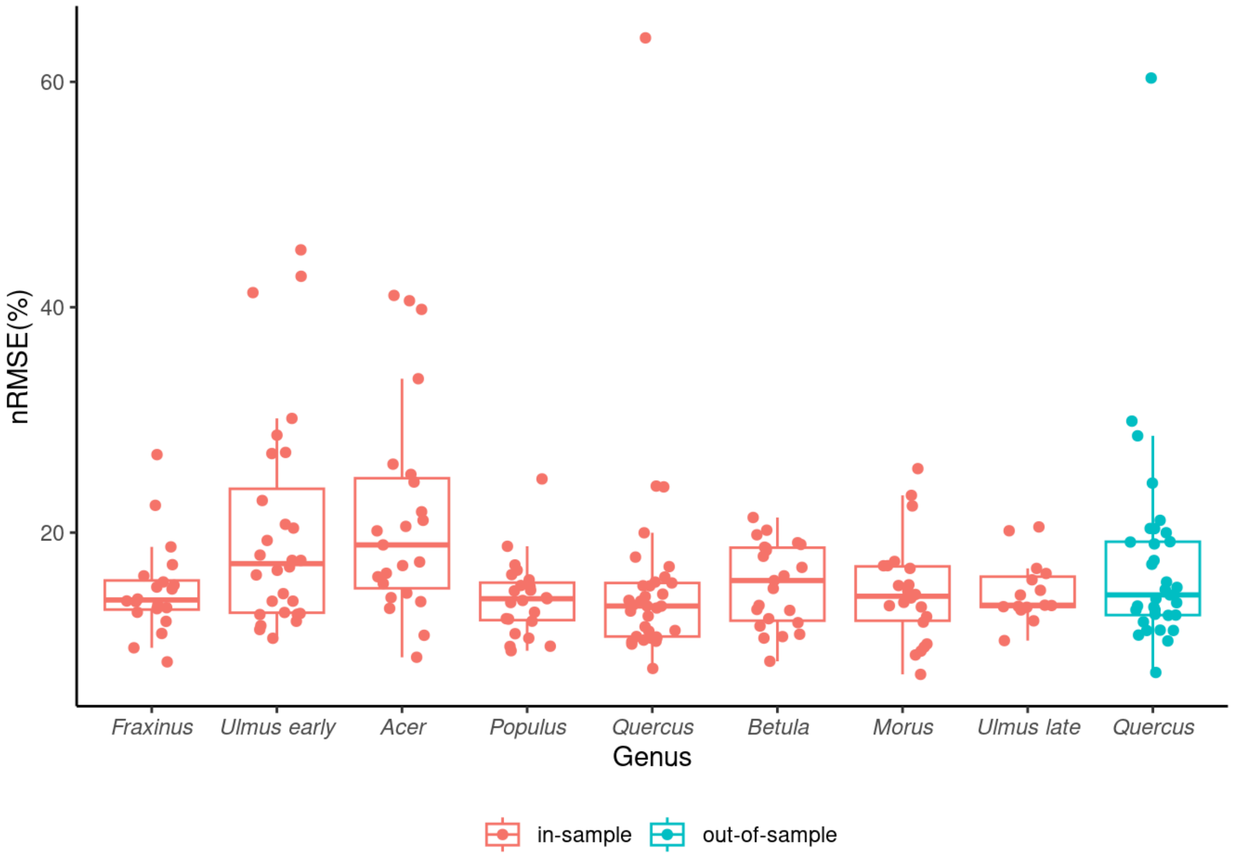


**Figure S10.** Accuracy of inferring pollen phenology with the PlanetScope method, both in-sample (fitting model with data from all cities) and out-of-sample (leave-one-out cross-validation), measured by normalized root mean square error (nRMSE, %).

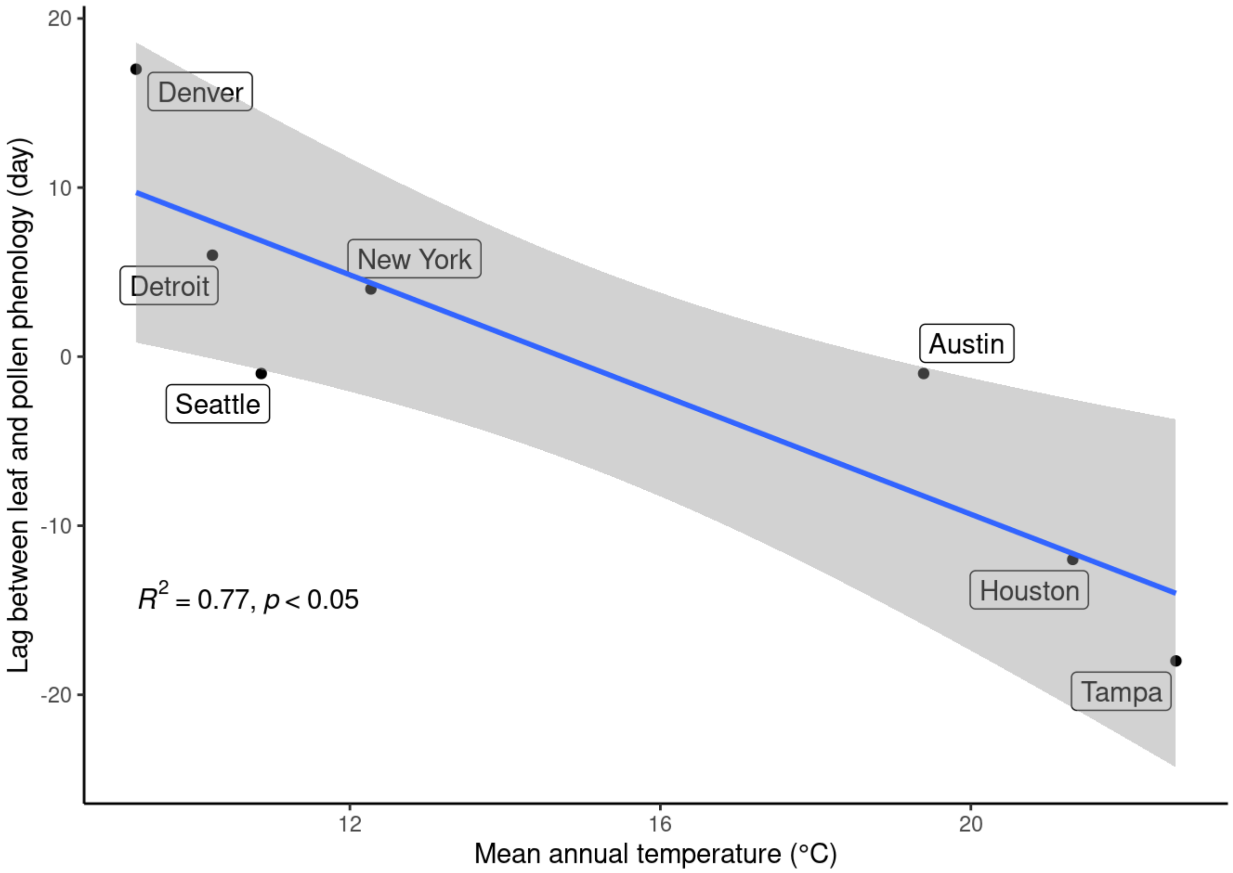


**Figure S11.** Linear regression between optimized lag between PlanetScope-derived leaf phenology and observed pollen phenology (day) and mean annual temperature (°C) of the city across seven cities studied. Blue line shows the linear regression line and gray ribbon shows standard errors of the fitted values.

**Table S1.** Source of street tree inventories for seven cities used to analyze the inference of pollen phenology from PlanetScope-derived vegetative phenology.

| City | Inventory source | URL |
|---|---|---|
| Austin | OpenTrees.org | https://opentrees.org/#pos=11/30.2623/-97.7426 |
| Detroit | Private request from City of Detroit, Michigan | |
| Denver | OpenTrees.org | https://opentrees.org/#pos=11/39.7273/-104.9455 |
| Houston | koordinates.com, mirrored from a previous dataset on mycity.houstontx.gov | https://koordinates.com/layer/25245-houston-texas-street-tree-inventory/ |
| New York | OpenTrees.org | https://opentrees.org/#pos=11/40.7056/-73.9764 |
| Seattle | OpenTrees.org | https://opentrees.org/#pos=11/47.6154/-122.33 |
| Tampa | TampaTreeMap | https://www.opentreemap.org/tampa/map/ |
